# Multiplexed 3D atlas of state transitions and immune interactions in colorectal cancer

**DOI:** 10.1101/2021.03.31.437984

**Authors:** Jia-Ren Lin, Shu Wang, Shannon Coy, Yu-An Chen, Clarence Yapp, Madison Tyler, Maulik K. Nariya, Cody N. Heiser, Ken S. Lau, Sandro Santagata, Peter K. Sorger

**Author notes:** These (first) authors contributed equally (Jia-Ren Lin, Shu Wang, Shannon Coy). These (senior) authors contributed equally (Sandro Santagata, Peter K. Sorger).

## Abstract

Advanced solid cancers are complex assemblies of tumor, immune, and stromal cells characterized by high intratumoral variation. We use highly multiplexed tissue imaging, 3D reconstruction, spatial statistics, and machine learning to identify cell types and states underlying morphological features of known diagnostic and prognostic significance in colorectal cancer. Quantitation of these features in high-plex marker space reveals recurrent transitions from one tumor morphology to the next, some of which are coincident with long-range gradients in the expression of oncogenes and epigenetic regulators. At the tumor invasive margin, where tumor, normal, and immune cells compete, T-cell suppression involves multiple cell types and 3D imaging shows that seemingly localized 2D features such as tertiary lymphoid structures are commonly interconnected and have graded molecular properties. Thus, while cancer genetics emphasizes the importance of discrete changes in tumor state, whole-specimen imaging reveals large-scale morphological and molecular gradients analogous to those in developing tissues.

## INTRODUCTION

Much of our knowledge of tumor microanatomy derives from 150 years of inspection of hematoxylin and eosin (H&E)-stained tissue sections, complemented for the last eighty years by immunohistochemistry (Coons et al., 1942). Histopathology has identified numerous recurrent features of tumors with diagnostic or prognostic significance (Amin et al., 2017), but classical methods often provide insufficient information for mechanistic studies and precision medicine. Spatial tumor atlases (Rozenblatt-Rosen et al., 2020) aim to build on this foundation and the current understanding of tumor genetics by collecting detailed molecular and morphological information on cells in a preserved 3D environment. The construction of such atlases is made possible by the recent development of highly-multiplexed tissue imaging methods (Angelo et al., 2014; Gerdes et al., 2013; Giesen et al., 2014; Goltsev et al., 2018; Lin et al., 2018; Saka et al., 2019; Schürch et al., 2020; Wagner et al., 2019) that yield subcellular resolution images of 10-80 antigens. When segmented and quantified, high-plex tissue images make it possible to identify cell types, assay proliferation, measure oncogene expression, and generate single-cell data that are a natural complement to scRNA-seq (Burger et al., 2021; Gaglia et al., 2022; Nirmal et al., 2022). Despite our increasingly deep knowledge about the genomic drivers of cancer – from oncogenic mutations to large-scale chromosomal rearrangements – we do not yet know how the spatial arrangement of the tumor microenvironment (TME) impacts pathogenesis; for instance, which feature types and spatial scales are relevant for mapping the 3D TME, how disease-associated histological features relate to molecular states, and whether morphological differences are discrete (like mutations) or continuous (like morphogen gradients found in development).

‘*Bottom-up*’ approaches to tissue atlas construction involve enumerating cell types, identifying cell-cell interactions, and generating local neighborhoods using spatial statistics. Such approaches leverage tools developed for the analysis of dissociated single cell data (e.g., mass cytometry (Bendall et al., 2011) and scRNA-seq (Luecken & Theis, 2019)). In contrast, “*top-down*” approaches involve annotating histopathologic features (histotypes) that have been demonstrated to associate with disease state or outcome (Amin et al., 2017) followed by computation on the multiplexed data to identify underlying molecular patterns. Histopathology has a long history of identifying striking spatial features in small cohorts that do not have prognostic or diagnostic value on follow-up, introducing a note of caution into ‘*bottom-up*’ analysis (Mazer et al., 2019; Voskuil, 2015). At the same time, discoveries arising from ‘t*op-down*’ analysis are strongly influenced by prior expectations. In this paper, we analyze colorectal cancer (CRC) using both approaches and compare the resulting insights.

Histological features of established significance in CRC include: (i) the degree of differentiation relative to normal epithelial structures based on tumor cell morphology (e.g., cell shape, nuclear size, etc.) and the organization of cellular neighborhoods (e.g., glandular organization, hypercellularity, etc.) (Fleming et al., 2012); (ii) the position and morphology of the invasive margin (Cianchi et al., 1997; Schürch et al., 2020) including the presence of “tumor buds,” small clusters of tumor cells surrounded by stroma (Lugli et al., 2017) which are correlated with poor outcomes (i.e., increased risk of local recurrence, metastasis, and cancer-related death) (A. C. Rogers et al., 2016); (iii) the extent of T-cell infiltration (Bruni et al., 2020) and the presence of peritumoral tertiary lymphoid structures (TLS) (organized aggregates of B and other immune cell types; (Di Caro et al., 2014)). In many cases, the origins and molecular basis of these histological features are not fully understood, although de-differentiation, including “stemness” (Aponte & Caicedo, 2017), epithelial-mesenchymal transition (EMT) (Kalluri & Weinberg, 2009), changes in nuclear mechanics (Uhler & Shivashankar, 2018), and similar processes, are likely involved. In the case of tumor budding, epigenetic changes, not specific mutations, have been shown to drive EMT (Centeno et al., 2017).

In this paper, we combined top-down and bottom-up analyses of high-plex CyCIF (Lin et al., 2018) and H&E images of CRC with single-cell sequencing and micro-region transcriptomics. We show that accurate assessment of disease-relevant tumor structures requires the statistical power of whole-slide imaging, not the small specimens found in tissue microarrays (TMAs); this typically corresponds to 10^5^ to 10^6^ cells per specimen, far more cells than are required for dissociative methods. Using 3D reconstruction of serial sections and supervised machine learning, we show that archetypical CRC histologic features are often graded and intermixed with morphological transitions and molecular gradients spanning 10^2^ or more cell diameters. Tumor budding also appears to be a graded phenotype, and budding cells, as classically defined, form an extreme example of a gradual molecular and morphological transition. Moreover, tumor buds, TLS, and several other structures are substantially larger than they appear in 2D: for example, B cell-rich TLS are interconnected communities of lymphocytes that can extend throughout large regions of the tumor. Thus, the TME is organized on spatial scales spanning 3-4 orders of magnitude, from subcellular organelles to cellular assemblies of hundreds of microns or more.

## RESULTS

### Overview of the specimens and data

Multiplexed CyCIF and H&E imaging were performed on 93 FFPE CRC human specimens spanning a wide variety of histologic and molecular subtypes (**Table S1**), which were imaged in three different formats, as illustrated in **Figure 1A**. Sample CRC1 (**Figures 1B-1E**) was subjected to 3D analysis: 106 serial sections were cut from an ∼1.7 × 1.7 cm piece of FFPE tissue and 22 H&E and 25 CyCIF images were collected, skipping some sections to increase the total dimension along the Z-axis. These images were then reconstructed in 3D and combined with scRNA-seq, and GeoMx transcriptomics (Zollinger et al., 2020) (**Figures 1A, S1A; Tables S1, S2**). Second, 16 additional samples (CRC2-17) were imaged in 2D as whole slides. Finally, CRC2-17 plus 77 additional tumors (CRC18-93) were imaged as part of a TMA (0.6 mm diameter cores; four cores per patient) (**Figure 1A**). In each case, CyCIF was performed using various combinations of 102 lineage-specific antibodies against epithelial, immune, and stromal cell populations and markers of cell cycle state, signaling pathway activity, and immune checkpoint expression (specific antibodies for the ‘main,’ ‘tumor-focused,’ and ‘immune-focused’ panels are listed in **Tables S3-S6**). MCMICRO software (Schapiro et al., 2022) was used to segment images, quantify fluorescence intensities on a per-cell basis, and assign cell types based on lineage-specific marker expression (**Figures 1C, S1B-S1C; Table S7**). Overall, ∼2 × 10^8^ segmented cells were identified in 75 whole-slide images using different combinations of antibodies (∼6TB of data) (Muhlich et al., 2022). All data are available for download via the HTAN Portal (see data access) and images of CRC1-17 are available for interactive online viewing without data or software download using MINERVA software (Hoffer et al., 2020; Rashid et al., 2022).

**Figure 1.**
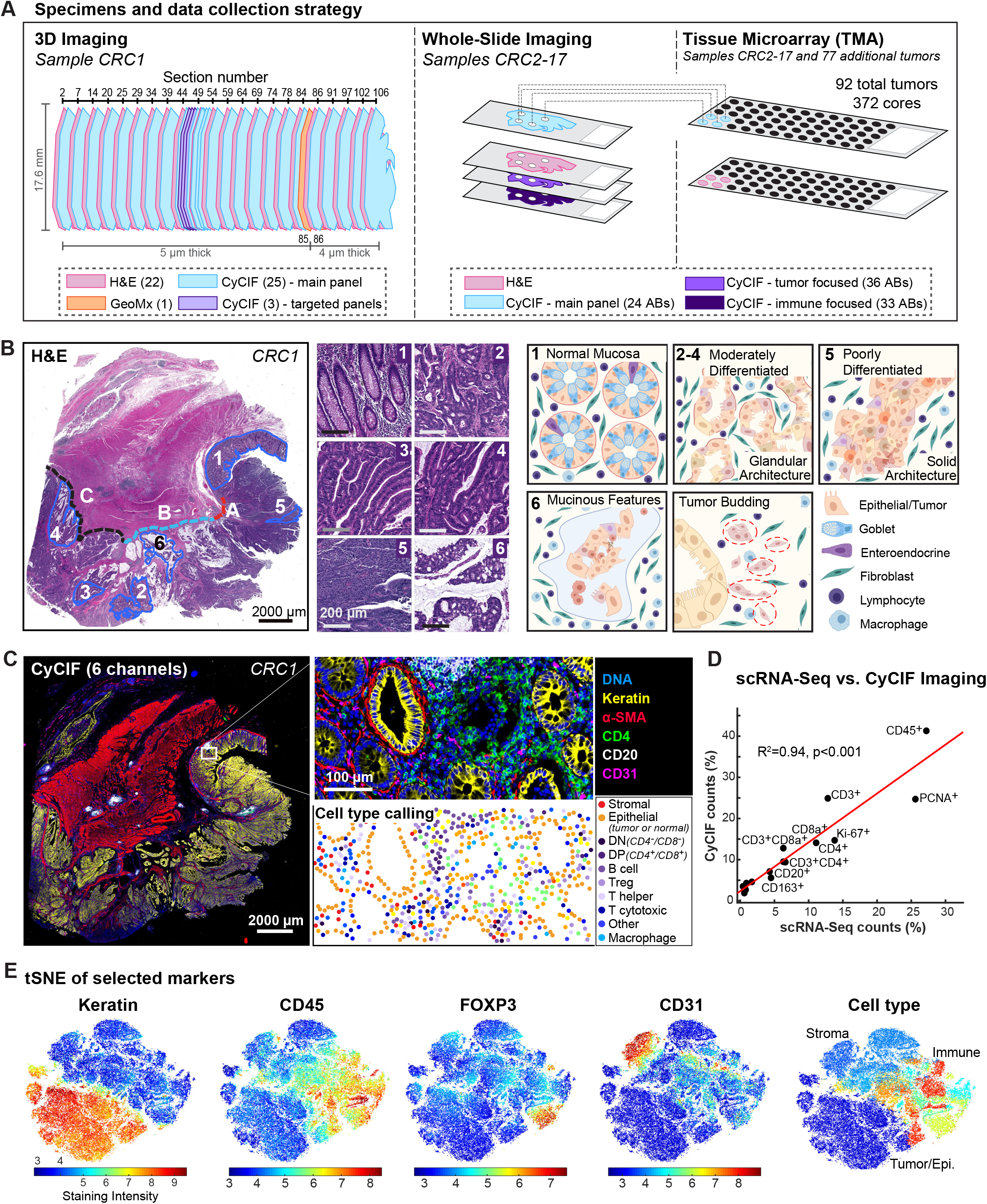
Overview of the data. **(A)** Overview of the data collection strategy for 93 FFPE CRC specimens available as a 3D stack, single whole-slides, and TMAs. Specimens were serially sectioned for H&E staining and CyCIF imaging (**Table S2**). For CRC1 (see **Table S1** for clinical information), 25 sections were stained with the main CyCIF antibody panel (**Table S3**), and 3 sections were stained with targeted panels (**Tables S4-S6**). Sixteen additional tumors (CRC2-17; **Table S1**) were imaged as whole-slides using the main and targeted antibody panels and also included in a TMA along with 77 additional CRC tumors (4 cores each; CRC18-93; **Table S1**). **(B)** Histopathologic annotation of H&E images into three invasive margins (A: budding margin, B: mucinous margin, C: pushing margin) and 6 different ROIs (*1: normal mucosa, 2: superficial (luminal) adenocarcinoma, 3: submucosal adenocarcinoma, 4: muscularis propria adenocarcinoma (deep invasive front), 5: solid adenocarcinoma, 6: mucinous adenocarcinoma)*. ROIs 2-4 exhibit a moderately differentiated appearance with glandular architecture, while ROIs 5-6 exhibit a poorly differentiated appearance with predominantly solid or cribriform architecture. Regions of tumor budding were also annotated. Schematic made with BioRender. **(C)** An example of a CyCIF whole-slide image (section 097) and cell-type assignment. Twenty-one different cell types from three main categories (tumor, stroma, and immune; **Table S7**) were defined and their locations mapped within the tumor section. **(D)** Comparison of cell-type percentages assessed via single cell RNA-sequencing (scRNA-seq) and CyCIF. **(E)** Dimensionality reduction of single-cell data by t-SNE (CRC1 section 097), color coded by staining intensity for the indicated marker. All markers were used to generate the t-SNE map while only random sampled 50,000 cells are shown in the plot. Cell-type plot (right) uses color code shown in **Figure S1C**.

Figure 1. shows images and single cell data for CRC1, a poorly differentiated stage IIIB BRAF^V600E^ adenocarcinoma (pT3N1bM0) (Weiser, 2018) with microsatellite instability (MSI-H) that arose in the cecum. This specimen was noteworthy for having complex histomorphology and an extended front invading into underlying smooth muscle (*muscularis propria*) and connective tissue. The front included a ‘budding invasive margin’ invading the submucosa adjacent to normal colonic mucosa (IM-A), a ‘mucinous invasive margin’ (IM-B), and a deep ‘pushing invasive margin’ (IM-C); the latter two regions invade the submucosa and muscularis (**Figure 1B**). t-SNE on the CyCIF data demonstrated a clear separation of cytokeratin-positive (CK^+^) epithelial cells (both normal and transformed) from CD31^+^ endothelial cells (primarily blood vessels), desmin^+^ stromal cells, and CD45^+^ immune cells (**Figures S1B-S1D; Table S8**). Immune cells could be further divided into biologically important classes such as CD8^+^PD1^+^ cytotoxic T cells (Tc), CD4^+^ helper T cells, CD20^+^ B cells, CD68^+^ and/or CD163^+^ macrophages, as well as discrete sub-categories such as CD4^+^FOXP3^+^ T regulatory cells (T_regs_) (**Table S7**). scRNA-seq was performed on ∼10^4^ cells from an adjacent (frozen) region of CRC1 (Chen et al., 2021) and the resulting estimated cell-type abundances exhibited a high degree of concordance with estimations from image data (R^2^ = 0.94; **Figures 1D-1E, S1E-S1F**), demonstrating the accuracy of the image segmentation and intensity quantifications.

### Impact of spatial correlation on statistical power

While having more single-cell data is preferable in principle, the effort required to collect 3D image stacks is substantially greater than single-section imaging; moreover, whereas a single whole-section image captures an individual patient’s data, an image of a TMA can contain specimens from >100 patients. For this reason, most high-plex tissue imaging papers to date focus on TMAs or – in the case of mass spectrometry-based imaging methods (MIBI, IMC) – on fields of view (FOVs) of ∼1 mm^2^. It is nonetheless well established that the minimum image dimension needed to accurately measure features within an image depends on the size of these features, which can be estimated from pixel-to-pixel correlation lengths (Rajaram et al., 2017). In CRC1-17, we observed correlation lengths ranging from ∼80 µm for CD31 positivity to ∼400 µm for keratin or CD20 positivity (**Figures 2A-2D, S2A**) and these length scales were directly related to observable and recurrent features of tumor morphology including capillaries for CD31^+^, sheets of tumor for CK^+^ cells, and TLS for CD20^+^ (**Figures 2C-2D**). Since these length scales are similar to those of most TMA cores, we used empirical and first-principles approaches to investigate the impact of sample size on the accuracy and precision of statistical analysis of 3D, 2D whole-slide, and TMA data.

**Figure 2.**
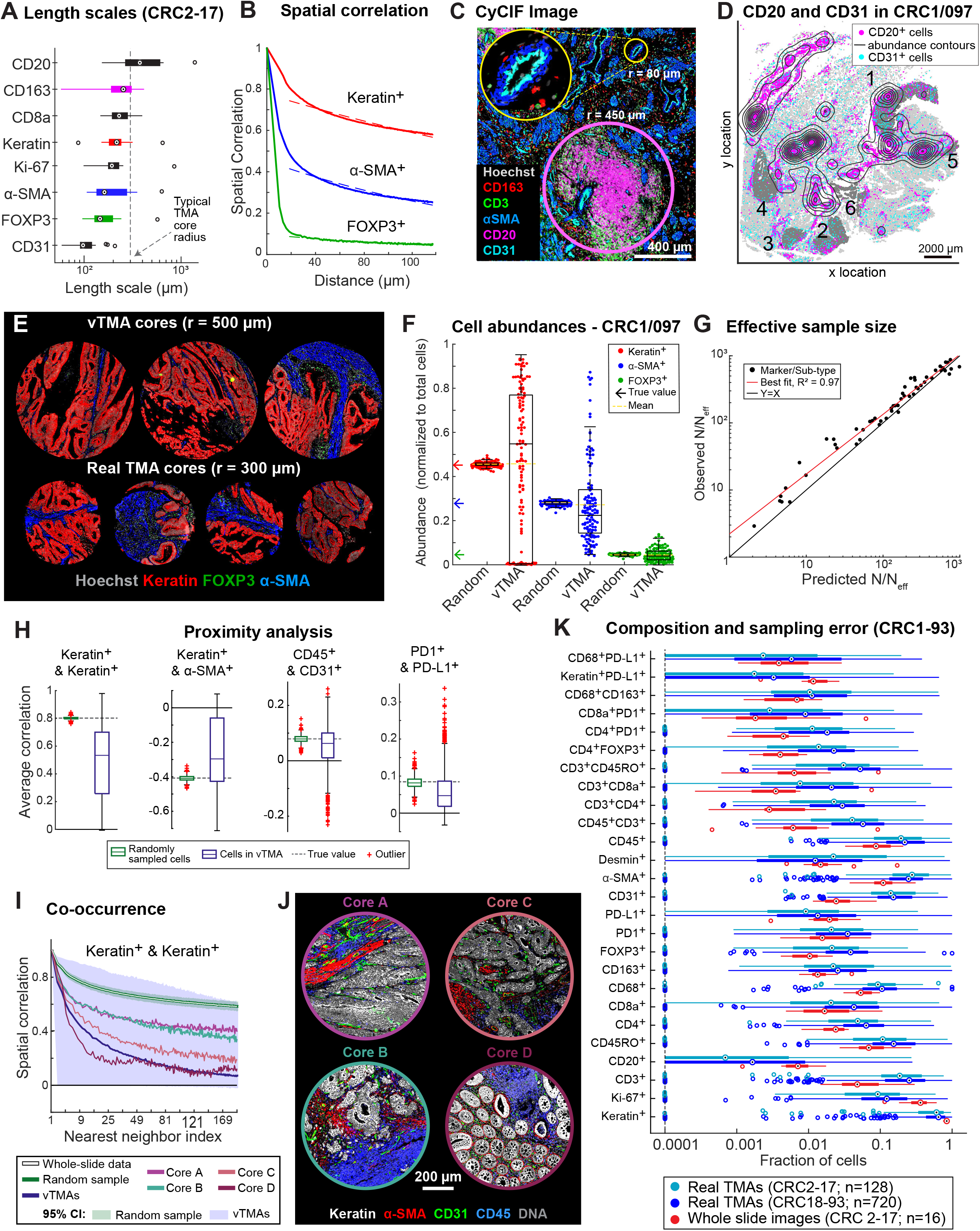
Spatial heterogeneity and estimation errors for regional sampling. **(A)** Length scales for select markers across CRC1-17. **(B)** Spatial correlations of binarized staining intensities for CK^+^ (red), α-SMA^+^ (blue), and FOXP3^+^ (green) cells, along with their exponential fits (dashed lines). **(C)** CyCIF image showing a CD20^+^ TLS (pink circle) and a CD31^+^ blood vessel (yellow circle and magnified in inset). (**D)** Spatial distribution of CD20^+^ cells (magenta dots and contours) and CD31^+^ cells (cyan dots); numbers 1-6 indicate annotated ROIs. **(E)** Virtual 1 mm TMA cores from CRC1 (section 097) and 0.6 mm cores from a real TMA of other CRC patients (CRC2-93). **(F)** Estimates of cell-type abundance using vTMA cores or random sampling. **(G)** Estimation error of vTMAs summarized by fold-reduction in effective sample size, N/N_eff_, for marker log-intensities and cell-type compositions. Observed error is compared to that predicted by accounting for exponential fits of spatial correlations in the Central Limit Theorem (R^2^ = 0.97, green); deviations (red) are attributable to some violation of model assumptions (e.g., deviation from exponential decay). **(H)** Correlation of select cell-type pairs amongst 10 nearest neighbors. **(I)** Correlation functions of CK^+^ cells as estimated from vTMAs or random sampling. Estimates from four cores are also shown. **(J)** Images of the four cores (A-D) highlighted in (I). **(K)** The fraction of various marker-positive cells across specimens CRC2-17 whole-slide or TMA data, or among TMAs from specimens CRC18-93. Box plot displays data points and 1^st^-3^rd^ quartiles, proportions <0.0001 are denoted as a single data point along the dotted line.

As an initial empirical approach, we generated a “virtual TMA” (vTMA) comprising 1 mm diameter FOVs subsampled from an image of CRC1 (section 097); each virtual core contained ∼10^3^ cells as compared to ∼5 × 10^5^ for a whole-slide CRC1 image. Sampling was performed so that the vTMA would primarily contain CK^+^ tumor or epithelial cells. CRC2-17 had been used, prior to the current work, to generate a real TMA (rTMA) in a pathology core, allowing us to confirm that vTMA and rTMA cores generated were similar (**Figure 2E**). When we computed the abundance of CK^+^ cells (cell count divided by the total cell number) in each vTMA core we found that it varied 20-fold from 5% to 95% whereas the true value determined by counting all cells in CRC1 section 097 was 45% (**Figure 2F**). Abundance estimates for α-SMA and FOXP3 positivity in vTMA cores were also imprecise, but to a lesser extent than for keratin positivity (**Figure 2F**). In contrast, when random samples of ∼10^3^ cells were drawn from the single cell data without regard to position in the specimen, the estimated abundance of CK^+^ cells was 45 ± ∼1%, a good estimate of the actual value (**Figure 2F**). Thus, imprecision associated with computing cell abundance from a vTMA arises only when spatial arrangements are preserved.

These findings can be explained in full from a theoretical perspective based on the Central Limit Theorem for correlated data (Lavrakas, 2008). The *effective* sample size (*N*_*eff*_) for correlated data is approximately related to the sample size *N* for “dissociated cells” (cells chosen at random without regard to position in an image or drawn from a dissociated cell preparation as in scRNA-seq or flow cytometry) via a simple scaling law (see Methods for the derivation):

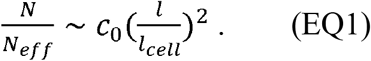

where *c*_0_ is the spatial correlation strength, *l* the length scale (e.g., ∼400 µm for CK) and average cell size *l*_*cell*_. We observed a good match between CyCIF data and theory (R^2^ = 0.97; **Figures 2G, S2B**) and a reduction in effective sample size (*N/N*_*eff*_) of 10-to 1,000-fold depending on the marker identity (median value ∼100). Thus, a 1 mm core containing ∼10^3^ spatially correlated cells constituted as few as 1 to 3 independent samples, which explains the high variance when cell abundance is estimated. We conclude that the analysis of TMA cores and other similarly small FOVs is an inadequate means to accurately determine a feature as simple as cell abundance simply because the sample is too small relative to the size of most features (we consider 2D v 3D sampling below).

Analysis of higher-order spatial features, such as cell proximity (**Figures 2H, S2C**) was also strongly impacted by spatial correlation. For example, vTMA data were much less precise than random sampling when computing the correlation of CK^+^ (tumor) cell frequency with neighboring α-SMA^+^ (stromal) cell frequency as a function of distance (compare blue and green in **Figure 2H**; note that distance is plotted as the number of neighboring cells, which is proportional to the square of the distance). The same was true when we searched for neighborhoods containing CD45^+^ immune cells and CD31^+^ endothelial cells that represent areas of perivascular inflammation. Inspection of underlying images showed that these differences related to variation of tissue morphologies and spatial arrangements (illustrated by four selected cores; **Figures 2I, 2J, S2D**).

To compare the magnitude of biological (patient-to-patient) variability with sampling error we computed cell abundances for single markers and biologically relevant marker combinations (e.g., CD68^+^PDL1^+^ macrophages) and observed a 3-to 10-fold variation from CRC2-17 (**Figure 2K**, red). However, inter-core variance from any single specimen obtained from rTMAs was substantially greater (**Figure 2K**, blue & teal). Only one measurement made from TMAs, Ki-67 positivity in CK^+^ cells, exhibited inter-patient variability (18-61%) greater than sampling error between cores (∼30%) (**Figures 2K, S2E-S2F**). Thus, imaging small fields of view causes sampling error to exceed true patient-to-patient variability in most cases. This error is sufficiently great that it can lead to false associations with patient outcome in Kaplan-Meier analysis (**Figures S2G-S2H**).

To determine whether 2D whole-slide images are an adequate approximation of a 3D specimens we computed cell abundances and spatial correlations for 24 Z-sections from CRC1 and compared this to patient-to-patient variability estimated from whole-slide images of specimens CRC2-17 (compare red and blue in **Figures S2I-S2J**). For all but a few markers, we found that variance between Z-sections was substantially smaller than patient-to-patient variability. The variances, when observed, were immediately interpretable as differences in tumor architecture along the Z axis. We conclude that 2D whole-slide imaging of a 3D specimen does not, in general, suffer from the same subsampling problem as TMAs or small fields of view; this too is consistent with theory about sampling under correlation. As we show below, however, some mesoscale features of tumors can only be detected in 3D datasets.

### Morphological and molecular gradients involving tumor phenotypes

To link high-plex image features to histological CRC features with well-established prognostic value, such as degree of tumor differentiation (well, moderate, poor), grade (low, high), subtype (mucinous, signet ring cell, etc.) (Weiser, 2018), two board-certified pathologists annotated regions of interest (ROI) from all 22 H&E sections of CRC1 and then transferred the annotations to adjacent CyCIF images for single-cell analysis. Annotations included normal colonic mucosa (ROI1); moderately differentiated invasive adenocarcinoma with glandular morphology involving the luminal surface (ROI2), submucosa (ROI3) or the *muscularis propria* at the deep invasive margin (ROI4); regions of poorly differentiated (high-grade) adenocarcinoma with solid and/or signet ring cell architecture (ROI5); and regions of invasive adenocarcinoma with prominent extracellular mucin pools (ROI6) (**Figure 1B**). A region with prominent tumor budding (TB) near margin IM-A was also annotated. Excluding muscle, CyCIF data showed that solid adenocarcinoma (RO15) had the highest proportion of CK^+^ tumor cells (∼70%), whereas adjacent normal epithelium (ROI1) had the fewest CK^+^ (∼25%) and the most stromal and immune cells.

To determine which molecular features correspond to each histology, we trained a k-nearest neighbor (kNN) classifier using molecular features (CyCIF intensities) on pathology labels. For simplicity, we consolidated the ROIs into four classes with half of the cells in each class used for training and half for validation. A different classifier was generated for each pair of CyCIF and H&E images for CRC1-17. Of note, the pathologist-labeled H&E data was rich in morphological context, but the CyCIF data comprised only cell positions (centroids) and integrated marker intensities, not morphological or neighborhood information. We observed high confidence predictions from the trained kNN classifier (Shannon entropy near zero) on the validation set (**Figures 3A, S3A**) showing that the classifier had encoded disease-relevant morphology using marker intensity alone. However, we found that no *single* molecular marker was unique to a specific ROI or tissue morphology implying that morphology is encoded in hyperdimensional intensity features.

**Figure 3.**
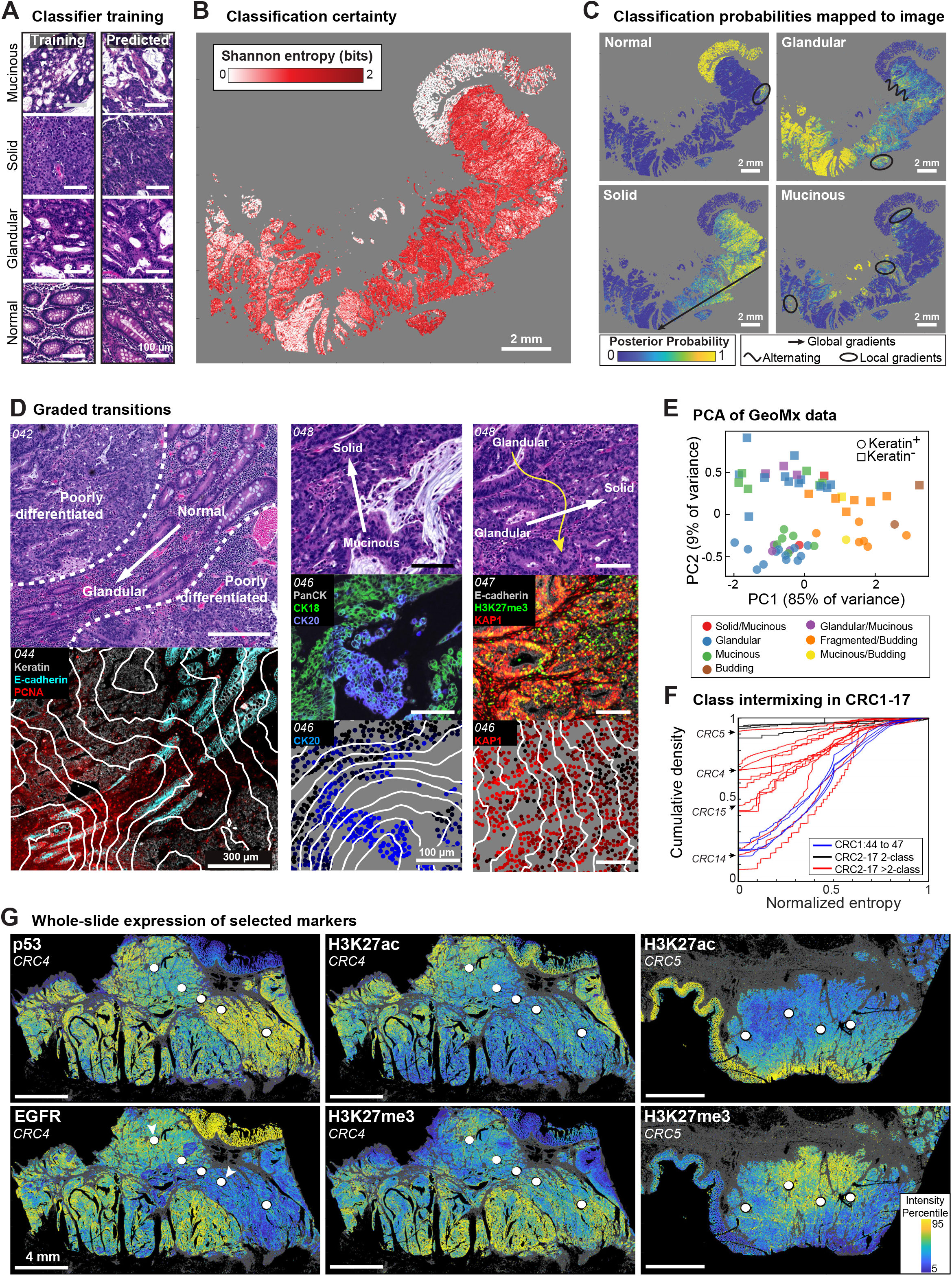
Correlation and prediction of morphologic and molecular tumor phenotypes. **(A)** Example ROIs corresponding to four different tumor morphologies (H&E) used for training (left column) and non-adjacent regions that were predicted with high confidence (right column). The kNN classifiers were trained and validated separately for each section to evaluate the reproducibility of the models. **(B)** Prediction confidence for assignment of kNN classes as measured by Shannon entropy. A value of 0 corresponds to perfect certainty. A value of 2 indicates random assignment (i.e., equal mixing of all classes). **(C)** Posterior probability that each CK^+^ cell belongs to normal epithelium or glandular, solid, or mucinous tumor classes. Annotation reflects classifier gradients and alternation corresponding to morphologic phenotype. **(D)** *Left:* Sample tumor region that transitions from normal to abnormal glandular features (H&E, top) coinciding with transition from E-cadherin expression to PCNA (CyCIF, bottom). Contours describe averaged local epithelial cell expression of PCNA. *Center and right:* Additional examples of transition regions, with H&E (top), CyCIF (middle), and quantified expression contours (bottom). **(E)** PCA of 31 spatially resolved GeoMx transcriptomics regions (analyzed areas indicated in **Figure S1A**). **(F)** Cumulative distribution of single-cell classification entropy of CRC1-17 rescaled to unit range. Patients with only two classes (black) had only normal epithelial and a tumor morphology class. Different CRC1 sections used different markers for classification. **(G)** Examples of marker gradients on the scale of whole tumor sections. White circles denote regions cored out during TMA construction.

Unexpectedly, kNN classifiers scored most regions of CRC1 outside of the annotated (training and validation) data as comprising a mixture of morphological classes (as quantified by the posterior probability) with transitions from one class to another. In many regions, Shannon entropy values approached two, demonstrating an equal mixture of all four classes (red in **Figures 3B, S3B**). This was not a limitation of the markers used for classification, because similar results were obtained with combinations of ∼100 antibodies used to stain sections 044 to 047 of CRC1 (**Figures S3C-S3D; Table S4**). When tumor regions with high Shannon entropy values were examined in H&E, we found that they corresponded to transitions between classical morphologies (**Figure 3D**). These transitions were not limited to a single part of the tumor but were observed multiple times in spatially separated areas on dimensions ranging from a few cell diameters (∼50 µm) to the whole image (∼1 cm) (**Figure 3C**) and included transitions from mucinous to glandular, mucinous to solid, and glandular to solid.

When we performed principal component analysis (PCA) on 31 spatially resolved transcriptomic microregions (using GeoMx microregion transcriptomics, with each microregion sorted into CK^+^ or CK^-^ cells) we also observed gradations in molecular state for both the tumor/epithelial (CK^+^; **Figure 3E**, circles) and immune/stromal (CK^-^; squares) compartments. In this case, principal component one (PC1; the dominant source of variance) correlated with histologic subtype and grade while PC2 correlated with epithelial vs. stromal compartment. In support of kNN models of CyCIF data, we observed a graded transition along PC1 from glandular/mucinous (low-grade) histologies to fragmented/budding (high-grade) histologies in both the epithelial/tumor and stromal/immune compartments. These findings serve to confirm the existence of graded state transitions at multiple locations in CRC1.

Across all 17 tumors, analysis of CyCIF data revealed intermixing of histologies to a greater or lesser extent with some tumors exhibiting contiguous blocks of a single morphology (e.g., CRC5) and intermixing similar to CRC1 in others (e.g., CRC14; **Figures 3F, S3B**). There was no obvious correlation between the degree of intermixing and MSI-H status (which promotes genome instability). We conclude that different and highly characteristic histological phenotypes routinely used for pathology grading and clinical planning in CRC are present in both discrete and intermixed forms, most likely due to epigenetic rather than genetic heterogeneity.

When we looked for patterns in CyCIF data we found that multiple markers exhibited intensity gradients that in some cases encompassed an entire tumor and in others coincided with local morphological gradients. Four examples are shown: a normal-glandular transition corresponding to E-cadherin and PCNA gradients that are inversely correlated (**Figure 3D**; left); a mucinous-solid transition coinciding with inversely correlated cytokeratin 20 and cytokeratin 18 gradients (**Figure 3D**; center); alternating glandular-solid transitions (**Figure 3D**; right, yellow curved arrow); and a glandular-solid transition coinciding with a transition in epigenomic regulators and modifications involving histone trimethylation (**Figure 3D**; right, white arrow). The histone acetylation (H3K27ac) vs. trimethylation (H3K27me3) marks are known to play complementary roles in transcriptional regulation (Zhao et al., 2021, p. 27), and we observed graded and anti-correlated expression on long length-scales in multiple tumors (e.g., CRC4, CRC5; **Figure 3G**), providing further evidence of organized epigenetic gradients in tumors. Graded expression of the tumor suppressor p53 and oncogene EGFR – two genes whose levels of expression play well established roles CRC biology – was also observed (**Figure 3G**). The white circles in **Figure 3G** are regions of tissue removed for rTMA construction (4 or 5 cores per specimen) based on the inspection of H&E images alone. It is immediately apparent that several sets of cores were inadvertently chosen to lie along a molecular gradient. Such variation between TMAs from a single specimen is often attributed to random heterogeneity rather than the large-scale structure we observe in whole-slide images. Chemical and physical gradients play essential roles in normal tissue development (Oudin & Weaver, 2016), but are less explored in tumors, perhaps because tumor genetics tends to focus on discrete differences (mutations).

### Tumor budding and molecular transitions at the deep invasive front

For diagnostic purposes, tumor buds are defined by the International Tumor Budding Consensus Conference (ITBCC) as clusters of ≤4 tumor cells surrounded by stroma and lying along the invasive front (Lugli et al., 2017), or, less commonly, the non-marginal ‘internal’ tumor mass (Lugli et al., 2011). Using ITBCC criteria a pathologist annotated buds in CRC1-17 and identified a total ∼7 × 10^3^ budding cells in 10 of 17 specimens examined (representing ∼0.01% of all tumor cells; **Figure 4A**, arrows and boxes highlight examples on H&E, yellow outlines on CyCIF images indicate segmented budding cells, **Figure S4A**). In CRC1, buds were largely confined to one ∼2.0 × 0.7 × 0.4 mm region of the invasive front (region IM-A, **Figure 1B**) near normal colonic epithelium and interspersed with T cells (**Figure 4B**). When we examined a 3D reconstruction of CRC1, we found that these “ITBCC buds” were frequently connected to each other and to the main tumor mass (**Figures 4C-4D, S4B**); buds as classically defined appeared to be predominantly cross-sectional views of these fibrillar structures (i.e., ‘bud-like’ structures rather than true buds). This observation is consistent with a previous 3D study of budding based on H&E images (Bronsert et al., 2014).

**Figure 4.**
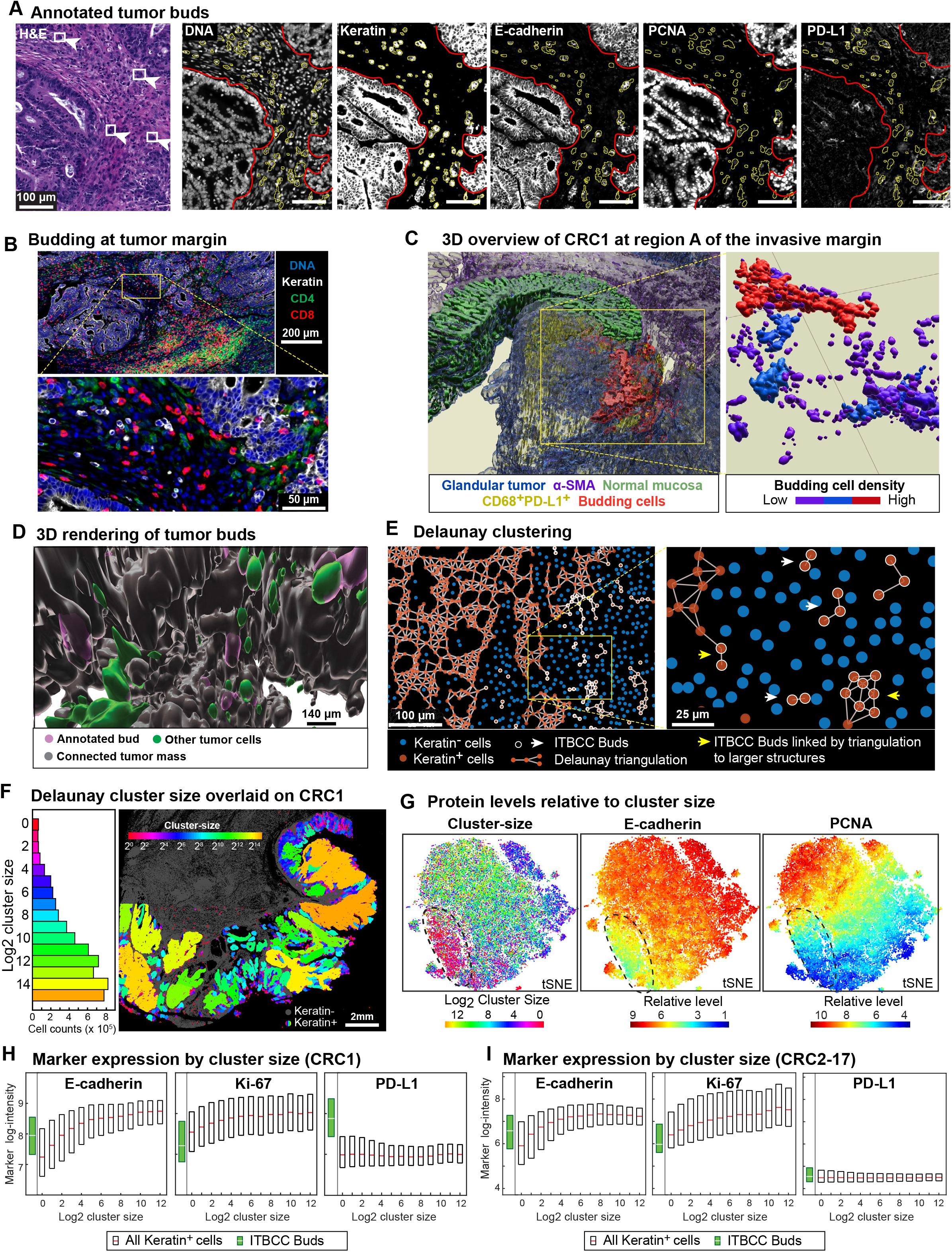
Tumor budding is a distributed phenomenon associated with graded molecular and morphologic transitions. **(A)** *Left:* H&E field of view (FOV) (CRC1, H&E section 096) from invasive margin A (IM-A, see **Figure 1B**) with a subset of budding cells indicated by boxes and arrowheads. *Right:* Corresponding CyCIF channels (CRC1, CyCIF section 097). Red outlines indicate the main tumor mass and yellow outlines the canonical tumor buds. **(B)** Different magnifications of the annotated budding region in CRC1 section 097. **(C)** 3D overview of CRC1 IM-A. *Left*: Surface renderings of glandular tumor (blue), α-SMA^+^ stroma (purple), normal mucosa (green), CD68^+^PDL1^+^ cells (yellow), and budding cells (red). *Right:* All annotated buds colored by budding cell density and showing interconnected fibril-like networks of budding cells. **(D)** 3D visualization of annotated buds (purple) relative to connected tumor mass (gray) and other cells with uncertain connectivity (green). Corresponding regions in 2D CyCIF images are in **Figure S4B. (E)** Delaunay clusters of CK^+^ cells in a local FOV of CRC1 section 097. CK^+^ cell neighborhoods are denoted by edges, along with CK^-^ cells (blue) and pathology annotated buds (white). **(F)** *Left:* Histogram of cluster-sizes (log2) across all 25 CRC1 sections. *Right:* Cluster sizes mapped onto CRC1 section 097. Image exaggerates size of single cells for visibility. **(G)** *Left:* t-SNE of cluster size. Color represents log2 cluster size and black outline denotes small clusters (including annotated buds). *Center and right:* t-SNE of CK^+^ cell expression of the indicated marker with color representing marker intensity. **(H)** Log-intensity of markers and their dependency on cluster-size in CRC1 tumor cells. Expression of annotated buds shown in green for reference. Box plots show 1^st^-3^rd^ quartiles; points beyond are not shown. Each box represents ∼10^5^-10^6^ cells. **(I)** Log-intensity of markers and their dependency on cluster-size for tumor cells in CRC2-17, as in **Figure 4H**.

To analyze these structures objectively, we used Delaunay triangulation (Delaunay, 1934), to annotate CK^+^ cells (i.e., tumor and normal epithelium) that were immediately adjacent to each other (**Figure 4E**). The smallest Delaunay clusters contained 1-4 contiguous cells surrounded by stroma and corresponded to ITBCC buds (**Figure 4F**; red) whereas the largest clusters contained >10^4^ cells and mapped to regions of poorly differentiated adenocarcinoma with solid architecture (which were almost entirely composed of tumor cells; yellow and orange). The widest range of cluster sizes was observed in differentiated regions with glandular architecture (**Figure 4F**; blue green). A key feature of tumor budding cells is that they express low levels of cell-to-cell adhesion proteins (e.g., E-cadherin, CD44, and Ep-CAM) (Gosens et al., 2007) and have a low proliferative index (Rubio, 2007, 2008). We confirmed that buds matching ITBCC criteria in our data had reduced expression of adhesion and proliferation markers (**Figure S4C**). Moreover, a t-SNE representation of all single cell data labeled by Delaunay cluster size showed that cells in the smallest Delaunay clusters expressed the lowest E-cadherin levels of all CK^+^ cells and that proliferation markers (e.g., PCNA) were also expressed at low levels (**Figure 4G**, circled region). However, tumors in our cohort did not contain a discrete population of E-cadherin/proliferation-low budding cells, instead, the expression of E-cadherin, Na-K ATPase, PCNA, and Ki-67 varied continuously with cluster size in CRC1 (**Figures 4H, S4D**) as well as other CRC tumors (**Figures 4I, S4E**).

Inspection of the underlying images (**Figures 5A-5B**) revealed that regions of cohesive glandular tumor (which was associated with large Delaunay cluster sizes and a PCNA^high^ state) often fragmented into fibrillar structures comprised of smaller clusters and a PCNA^low^ state. At the terminal tips of these fibrillar structures were ‘bud-like’ structures exhibiting the lowest PCNA expression and surrounded by stroma (**Figure 5A**) or mucin (**Figure 5B**). Analogous transitions between tumor masses and small Delaunay clusters were observed throughout the tumor both at the invasive front (IM-A in CRC1), in mucinous spaces (IM-B), and along the luminal surface of the tumor in regions corresponding to discohesive growth with focal signet ring cell morphology (ROI5, **Figure 1B**) (Sung et al., 2008). The small Delaunay clusters found in mucin pools were not distinguishable in size or expression (of cohesive and proliferation markers) from buds as classically defined (**Figures 4I, S4E**) even though the ITBCC definition encompasses only clusters in fibrous stroma. PCA of GeoMx RNA expression data (**Figure 3E**) confirmed that regions with ITBCC buds (brown dots), fragmented tumor and budding (orange), and budding into mucinous spaces (yellow) were similar to each other and distinct from other tumor morphologies (**Figure 3E**). Moreover, all three bud-like morphologies expressed elevated levels of genes in the EMT Hallmark gene set (GSEA M5930; **Figure 5C**, orange, yellow, brown) consistent with the idea that a loss of cell cohesion occurs frequently across tumors, is associated with an EMT-like process, and may be driven by a similar epigenetic program (Centeno et al., 2017). In 2D views, mucin surrounding many bud-like structures in CRC1 is found in pools of many different sizes that are apparently isolated from each other (mucins are large glycoproteins that protect the gastrointestinal epithelium; **Figure 5D** arrowheads) (Bresalier, 2002). However, when we used the CRC1 3D reconstruction to map these pools, we found that they were frequently continuous with each other and with the colonic lumen, up to 1cm away; in CRC1 this is most prominent in the central region involving invasive margin IM-B (**Figure 5E**).

**Figure 5.**
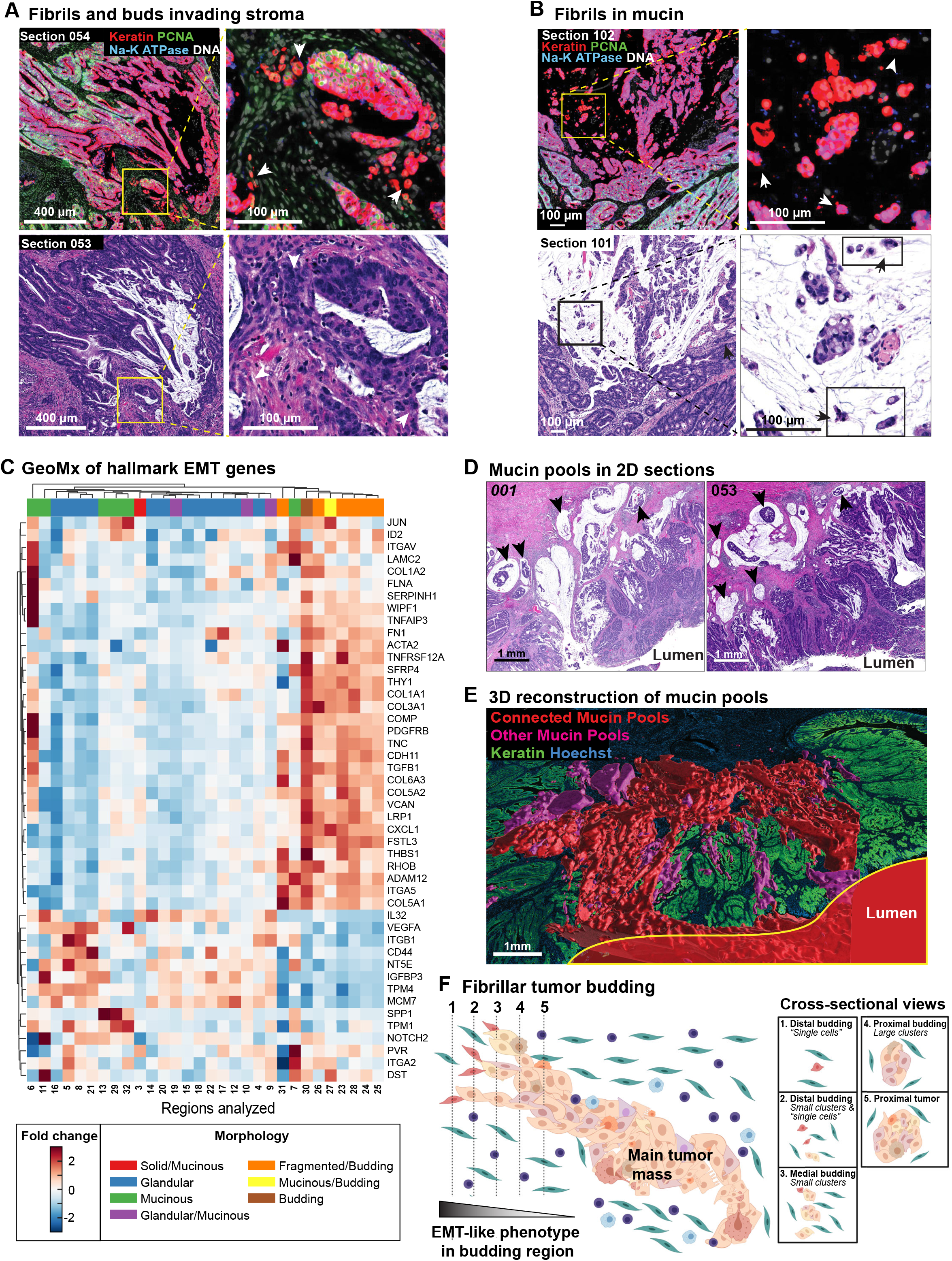
Small, isolated tumor and mucin structures in 2D are large, connected networks in 3D. **(A)** Example of transition from main tumor mass into fibrils and ‘bud-like’ cells in the stroma in CyCIF (*top*) and H&E (*bottom*). There is gradual loss of Na-K ATPase and PCNA from the main tumor mass to the tips of fibrils with decreasing cluster-size (budding cells appear red on CyCIF, arrowheads). Image is oversaturated to make hues more visible. **(B)** Analogous budding structures in mucinous tumor regions, with fibrils and budding cells (arrowheads) extending into mucin pools rather than stroma. **(C)** Heat map of GeoMx data for selected EMT hallmark genes. Each column corresponds to an analyzed region from one tissue section, as described in **Figure S1A**. Morphology corresponding to each region is indicated. **(D)** Two exemplar H&E FOVs from different regions of the reconstructed mucin structure with mucin pools that appear isolated in 2D sections (arrowheads). **(E)** Connectivity of mucin pools across serial sections. Largest contiguous mucin pool structure (red) extends into the lumen (lumen surface outlined in yellow). Image is mirrored along Z relative to **Figure 1B** to better visualize details. **(F)** Schematic diagram depicting serial sectioning through fibrils of budding cells at the tumor invasive margin, illustrating how a large contiguous 3D structure may appear as isolated cells or small clusters in 2D sections. Made with BioRender.

Putting these data together, we conclude that EMT-like transitions and tumor budding in CRC is characterized not by the formation of isolated spheres of cells, as first described by Weinberg and colleagues in tissue culture (Mani et al., 2008), but instead by the formation of large fibrillar structures that appear to be small buds when viewed in cross-section at their distal tips. Fibrils can invade into a variety of different environments including stroma and mucin (which itself consists of large inter-connected mucin-filled structures rather than isolated pools) and we speculate that their formation is driven by a gradual (not abrupt) breakdown in cell adhesion associated with a graded EMT-like transition (**Figure 5F**).

### Networks of tertiary lymphoid structures and their composition

Anti-tumor immunity involves innate as well as adaptive mechanisms that mediate the expansion and activation of cytotoxic T cells and the production of antibodies by B cells (plasma cells). Adaptive immunity occurs within secondary lymph organs (Peyer’s patches in the colonic mucosa) (Schumacher & Thommen, 2022) as well as tertiary lymphoid structures (TLS), which develop in non-lymphoid tissues such as tumors and other sites of chronic inflammation. The formation, organization, and functions of TLS are under active investigation, but their presence is known to be associated with good prognosis and immune checkpoint inhibitor (ICI) responsiveness (Cabrita et al., 2020; Helmink et al., 2020). Pathology inspection of 47 individual sections of CRC1 (22 H&E and 25 CyCIF) identified over 900 distinct SLO and TLS domains in 2D (**Figures 6A, S5A**). However, following 3D registration and segmentation, we found that many of these domains were connected in larger 3D structures; for example, seven large networks (**Figure 6B**; 3D rendering view) each spanning >12 sections and several mm laterally, could be assembled from 20-200 individual 2D domains each (the final assembly included 133 additional smaller SLO/TLS networks; **Figures 6C, S5B**). The large tertiary lymphoid structure networks (TLSNs) were found along the invasive fronts (networks A, B, D), inside tumor (F, G), or in layers of the *muscularis* (E) or *subserosa* (C; the subserosa is peri-colonic fibroadipose tissue external to the muscularis). To study the cellular composition of TLSNs, we performed K-means clustering on CyCIF intensity data (with k = 7 to match the number of large networks, **Figure 6D**) and recovered clusters with the properties of SLOs (cluster 3) that were found near normal mucosa (as expected for Peyer’s patches) or typical TLS-like lymphoid-aggregates within the tumor itself (cluster 1, **Figures 6E-6F, S5C-S5D**). TLS undergo maturation and are expected to differ from one another, but when we mapped marker expression clusters onto the physical organization of TLSNs, we found that some were relatively homogenous, containing cells from one expression cluster whereas others were heterogenous. For example, TLSN-C, which was predominantly located in the *subserosa*, was >96% composed of expression cluster 7, which showed a marked predominance of CD45^+^CD20^+^ B cells with little enrichment of other populations; TLSN-F, which was found immediately adjacent to the region of tumor budding, was 95% comprised of cluster 6 which was defined by a more heterogeneous collection of immune lineages including B cells, numerous PD1^+^ cytotoxic T cells, FOXP3^+^ T_regs_, and PDL1^+^ myeloid cells. In contrast, other TLSN-A, -B, or -D contained mixtures of expression clusters (**Figures 6E, S5C**).

**Figure 6.**
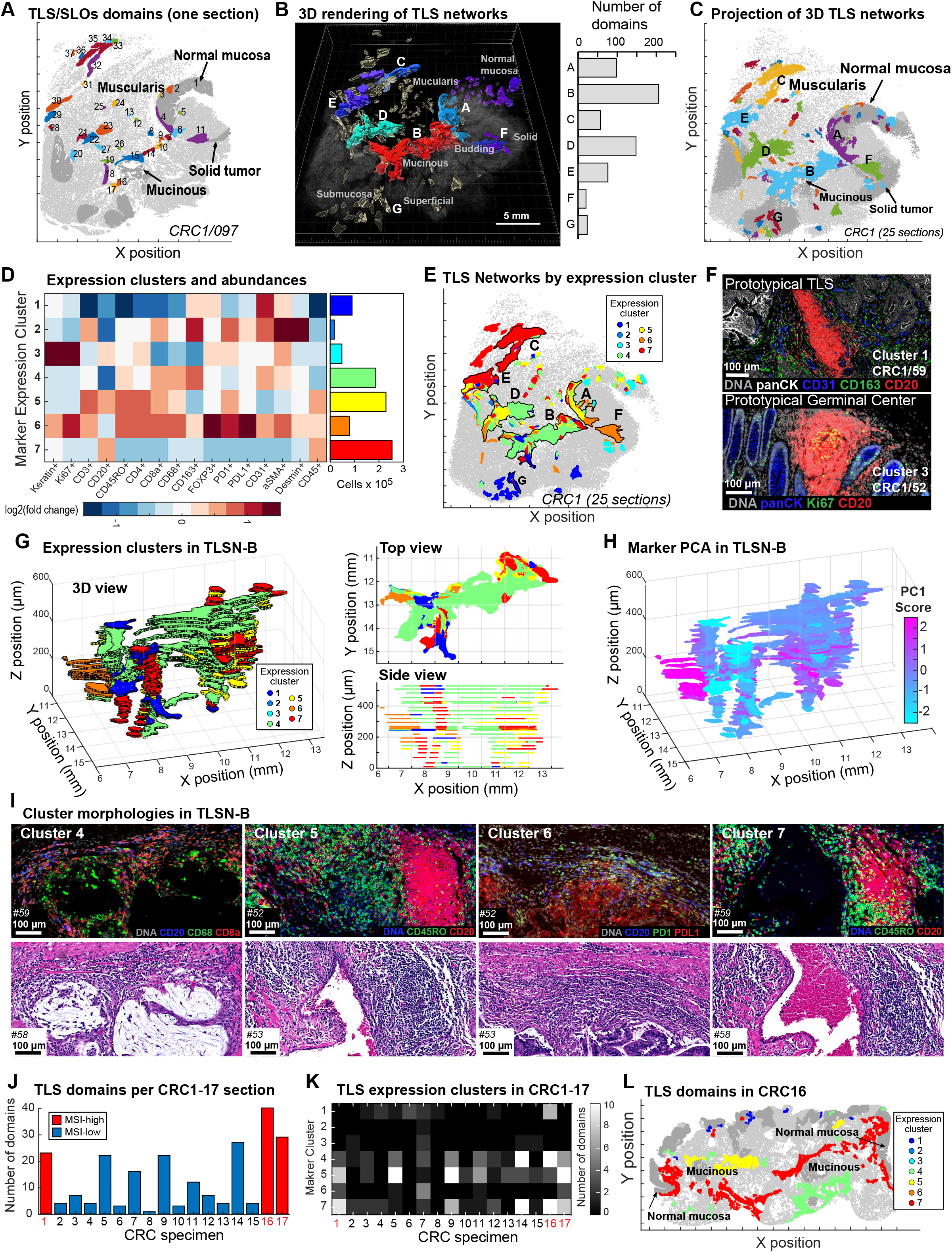
3D TLS structure and cell compositions. **(A)** 2D TLS domains in CRC1, section 097. Numbers indicate the individual TLS/SLO domains identified in this one section. **(B)** 3D rendering view of TLS networks (TLSNs) from CRC1. The 7 largest TLS networks are labeled A-G. Histogram shows the number of individual TLS identified in 2D sections from each TLS network (A-G). **(C)** 3D TLS networks projected onto XY surface. **(D)** Clustering of TLS domains by kNN (left panel) and number of domains in each cluster (right panel). **(E)** TLS cluster distribution in CRC1; 7 largest TLSNs are outlined and labeled. **(F)** Example CyCIF images of TLS clusters 1 and 3. **(G)** *Left:* 3D view of TLSN-B from CRC1 with each TLS domain colored by cluster. This 3D view is in the same orientation as TLSN-B shown in panel E and in the top view (shown in right upper panel). *Right*: Cross-sectional views of XY (top) and XZ (bottom) show TLS domains in TLSN-B. **(H)** 3D view of TLSN-B, colored by principal component 1 (PC1). **(I)** Example CyCIF and H&E images of TLS clusters 4, 5, 6, and 7. **(J)** TLS domain counts in CRC1-17 (section 097 was used for CRC1). **(K)** Heatmap of TLS clusters from CRC1-17. **(L)** 2D TLS domains of CRC16, colored by TLS clusters.

To study an intermixed TLSN in greater detail, we projected marker clusters onto a 3D rendering of TLSN-B (**Figure 6G**), which had been assembled from the greatest number of individual 2D domains (206) and spanned all sections of CRC1 (**Figures 6B, S5B**). We found enrichment of myeloid cells (CD68^+^CD163^+^; cluster 4, green) on the mucinous side of TLSN-B, with enrichment of T-cell (CD3^+^, CD45RO^+^, CD4^+^; cluster 5, yellow) and B-cell (CD20^+^CD45^+^; cluster 7, red) clusters intermixed along the stromal side (**Figure 6G**). Inspection of corresponding H&E images revealed numerous discrete B cell aggregates with associated T cells in clusters 4 to 7 with states distinguished by the relative abundance of different cell types (**Figure 6I**). The impression of graded composition was confirmed when we performed PCA on marker intensities and mapped principal component scores onto the TLSN-B structure (**Figures 6H, S5E**). This representation of the data emphasized the gradations in composition found within a single network.

To extend this analysis, we superimposed the marker-based clustering from CRC1 onto CRC2-17 (**Figure S5F**); we found that the prevalence of individual marker clusters varied from tumor to tumor but was similar for CRC1 and CRC2-17 in aggregate (**Figures 6J, 6K**). Like CRC1, CRC16 and 17 are MSI-H tumors with rich TLS networks that appear large and connected even in 2D. Moreover, in CRC16 the area surrounding mucin pools and TLS were enriched in cells from marker clusters 4, 5 and 7 – as in CRC1 (**Figure 6L**) From these data, we conclude that CRC1 is a reasonable exemplar of our overall cohort and that preliminary conclusions can therefore be drawn from our single 3D reconstruction. These are that (i) TLS can form interconnected networks rather than the isolated structures observed in 2D sections, (ii) TLS networks within a single tumor can differ from one another significantly with respect to the proportions of different immune lineages, and (iii) the cell types and functional markers within a single large TLS network can vary considerably from one region to the next, and much of this variation appears graded, implying intra-TLS patterning and communication.

### Immune profiling of the invasive margin

The immune response at the tumor margin strongly influences disease progression and ICI responsiveness (Paijens et al., 2021). Among the three morphologies found at the CRC1 invasive margin, IM-A, the region with tumor budding and poorly differentiated morphology, had the greatest density of immune cells (**Figure 7A**) but was also strongly immunosuppressive, with abundant CD4^+^FOXP3^+^ T_regs_ partially-localized with CD8^+^ cytotoxic T cells along tumor margins (**Figure 7B**). While PDL1^+^ cells were found inside the tumor and the stroma (**Figure 7C**), the interaction between PDL1^+^ and PD1^+^cells was enriched at the budding interface cells (**Figure 7D**). IM-B exhibited the least immune cell infiltration, consistent with a role for mucins in immune evasion or sequestration (Bhatia et al., 2019). IM-C was rich in T_regs_ but had very few PDL1^+^ cells as compared to IM-A (**Figures 7C, 7D**). To explore the connection between the tumor margin morphologies and molecular properties systematically, we used using Latent Dirichlet Allocation (LDA), a probabilistic modeling method that reduces complex structures into distinct component communities (“topics”) while accounting for uncertainty and missing data (Blei et al., 2003; Jackson et al., 2020; Valle et al., 2014). We annotated invasive margins in CRC1-17 for i) infiltration with tumor budding, ii) deepest invasion, and iii) all other morphologies (mucinous fronts were too infrequent to represent their own category) and then performed LDA on CyCIF data (33-plex immune panel; **Figure S6A**) (Nirmal et al., 2022). We found that LDA topics had significantly different frequencies in different regions of the invasive margin (**Figures 7E, S6B-S6C**). Margins with tumor budding were significantly associated with CD4^+^ and CD8^+^ T cells (**Figure 7E**, topic 1), the deep invasive front with tumor cell proliferation (Ki-67 positivity in CK^+^ cells; topic 9), and the remainder of the front with podoplanin positivity (PDPN^+^; topic 7). PDPN is a short transmembrane protein widely expressed in cancer cells and cancer-associated fibroblast that is implicated in cell migration, invasion, and metastasis (Krishnan et al., 2018). Fibroblasts secrete abundant cytokines and growth factors, potentially explaining the activation of signal transduction (i.e., phosphotyrosine - pTyr - and phospho-SRC positivity; topic 10) along this portion of the tumor margin. In contrast, myeloid cells were ubiquitous, and their frequency (topics 5 and 12) did not significantly associate with any specific margin morphology. We conclude that morphologically distinguishable domains of the CRC invasive margin have differing levels of tumor cell proliferation (low in buds and high in deep invasive margins), activation (pTyr), and immune suppression.

**Figure 7.**
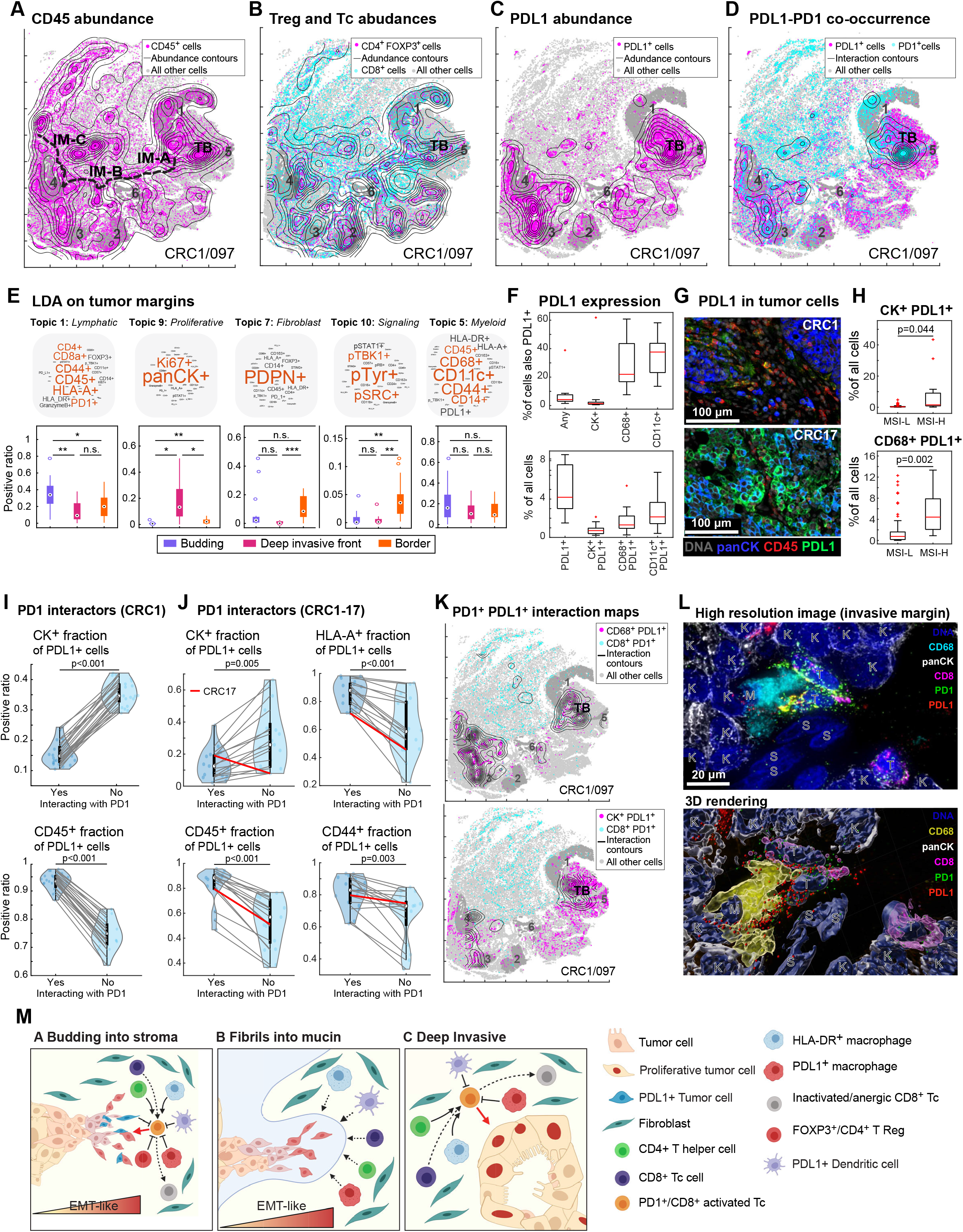
Immune landscape of CRC and its invasive margins. Abundance and distribution of **(A)** CD45^+^, **(B)** CD4^+^FOXP3^+^ (T_reg_) and CD8^+^ (Tc), and **(C)** PDL1^+^ cells with ROI numbers indicated; TB denotes region enriched for tumor budding; in panel **A**, the three regions of the invasive front are labelled IM-A, IM-B, and IM-C. **(D)** Co-occurrence of PDL1^+^ and PD1^+^ using a distance cutoff of 20 μm. Panels A-C and K depict CRC1 section 097. **(E)** LDA topics and their relative abundancies along the tumor margin. **(F)** PDL1 expression in indicated cell types. The top panel represents the relative fractions of PDL1^+^ cells over indicated populations, while the bottom panel shows the absolute fractions of PDL1^+^ or double marker positive cells. **(G)** Representative images of PDL1^+^CK^+^ cells in CRC1 (top panel) and CRC17 (lower panel). **(H)** Plot of PDL1^+^CK^+^ (top panel) or PDL1^+^CD68^+^ cell fractions in MSI-H or MSI-L samples from TMA data (CRC2-93). **(I)** Characterization of PDL1:PD1 interaction in CRC1 (all 25 sections). PDL1^+^ cells were binned into two subsets, one with PD1^+^ cells in proximity (20 µm cutoff) and one without. The fractions of CK^+^ (top panel) and CD45^+^ (bottom panel) in these two bins are plotted; P-values from pairwise t-test are shown (n = 25). **(J)** Characterization of PDL1:PD1 interaction in CRC1-17 performed as in panel I (n = 17). **(K)** Co-occurrence maps using a distance cutoff of 20 μm and cell types shown. **(L)** High-resolution 3D imaging of PDL1:PD1 interaction among tumor and myeloid cells. *Top panel*: maximum intensity projections. *Bottom panel*: 3D rendering from Imaris software. **(M)** Schematic illustrating tumor-immune interactions at different types of invasive margins.

With respect to immunosuppression, the distribution of PD1^+^ and PDL1^+^ cells is of particular interest because this interaction can be targeted therapeutically in CRC (André et al., 2022). Across CRC1-17, the fraction of PD1^+^cells varied 4-fold (from 3-12% of all cells) and these cells were >80% CD4^+^ or CD8^+^ T cells (**Figures S6D, S7**). The fraction of PDL1^+^ cells varied 12-fold (3-40%) (**Figure S6E**) and was correlated with the number of PD1^+^ cells (r = 0.52, p = 0.034; Z test). While a small minority (1-5%) of tumor cells expressed PDL1, the cells most likely to be PDL1^+^ were CD68^+^ (14-51% positive) and CD11c^+^ myeloid cells (10-88% positive); PDL1^+^ myeloid cells were also ∼6.5-fold more abundant on average than PDL1^+^ tumor cells (**Figures 7F, S6E**). The sole exception to this rule was CRC17 in which >40% of all tumor cells were strongly PDL1 positive; this tumor was also high-grade with extensive necrosis and uniformly poorly differentiated solid architecture and t-SNE showed it to be a clear outlier with respect to composition (**Figures 7G; S7A-S7C**). Immunotherapy is indicated for MSI-H CRCs because they are highly immunogenic (Boland & Goel, 2010); we found that MSI-H tumors in our cohort (n = 16 out of 93; see methods) had 5-fold more PDL1^+^ tumor cells and 6-fold more PDL1^+^ myeloid cells on average than MSI-L tumors (p = 0.044 and 0.002 two-side t-test, **Figure 7H**), but the latter still outnumbered the former ∼4-fold. Moreover, ∼80% of MSI-H tumors had more PDL1^+^ myeloid cells than the average MSI-L tumor (**Figure 7H**). Across the CRC cohort we found that single positive CD68^+^CD11c^-^ and CD68^-^CD11c^+^ as well as double positive CD68^+^CD11c^+^ cells were commonly PDL1^+^, although this fraction and the relative abundance of each myeloid subset varied several fold (**Figures S6F-S6G**), We do not have the markers in our panels to more precisely subtype PDL1^+^ myeloid populations across the CRC cohort but our interpretation is that they include variable proportions of macrophages, dendritic cells, and other mononuclear phagocytes.

Functionally, it is not simply the prevalence of PDL1^+^ cells that is relevant for T-cell suppression, but also which cells are in spatial proximity to allow for PDL1:PD1 binding. To study this, we performed proximity analysis using a 20 µm cutoff and found that, across 24 CRC1 sections, cells interacting with PD1^+^ cells were strongly enriched for CD45 positivity and depleted for CK positivity (p<0.001 pairwise t-test, two-sided), showing that PD1^+^ T cells more commonly interact with PDL1^+^ immune cells than tumor cells in CRC1. This was also true of CRC2-16 with CRC17 representing the sole exception (**Figure 7J**, red lines). Cells interacting with PD1^+^ cells were also significantly more likely to be positive for the CD44 adhesion receptor (Senbanjo & Chellaiah, 2017) and the HLA-A major histocompatibility antigen than non-interacting cells. Co-localization of CD68^+^PDL1^+^ myeloid cells with PD1^+^CD8^+^ T cells was also confirmed by co-occurrence mapping in CRC1 (**Figure 7K**, upper panel). Finally, high resolution (∼220 nm laterally) optical sectioning 12-plex CyCIF provided direct evidence of PDL1^+^ myeloid cells synapsing with PD1^+^ T cells at the tumor margin: we observed multiple examples of co-localization and polarization of PD1 and PDL1 on the membranes of adjacent cells, consistent with formation of functional cell-cell interactions (**Figure 7L**). We conclude that immunosuppression of PD1^+^ T cells in our CRC cohort most commonly involves PDL1^+^ myeloid, not PDL1-expressing tumor cells. Nevertheless, PDL1-expressing tumor cells may still be involved in immune suppression in some tumors. In CRC1 for example, greater >85% of interactions of PD1^+^ T cells with PDL1-expressing cells are myeloid in origin, but the 3% of tumor cells are that are PDL1^+^ are concentrated at the budding margin in close proximity to T cells (**Figure 7K**, lower panel; summary schematic **Figure 7M**).

## DISCUSSION

Understanding intra-tumor heterogeneity (ITH) is widely regarded as essential for improving our knowledge of tumor initiation and progression and ultimately for optimizing diagnosis and therapy (Marusyk et al., 2012). The image-based single cell analysis described in this paper supports two broad conclusions about the nature and organization of the ITH in CRC. First, our data show that molecular states (protein markers) and tissue morphologies (histotypes) are often graded, with transitions between phenotypes spanning spatial scales from a few cell diameters to many millimeters. For example, gradients in the epigenetic markers H3K27me3 and H3K27ac can span several centimeters along an entire tissue specimen. These markers play complementary roles in regulating transcription (Zhao et al., 2021, p. 27), and we find that their levels are commonly anti-correlated. In other cases, changes in cellular phenotypes are graded or recur in a semi-periodic manner, reminiscent of the “reaction-diffusion” gradients of morphogens observed in embryonic development (Turing, 1952) and also observed by fixed and intravital imaging in the mouse (Kondo et al., 2021) and in frozen human tissue by mass spectrometry (Randall et al., 2020). Second, cell-cell interactions most commonly studied at a local level are often organized into large and interconnected structures that are substantially larger than inspection of 2D sections suggests. These structures include: (i) the 1-4 cell tumor buds that are cross-sectional views of fibrillar structures (Bronsert et al., 2014) characterized by progressively lower expression of cell adhesion and proliferation markers as they narrow in diameter along the proximal-to-distal axis; (ii) intertumoral mucin pools that are surrounded by tumor in 2D but comprise 3D networks connected in some cases to the intestinal lumen and its microbiome; (iii) TLS, which are strongly implicated in anti-tumor immunity (Edin et al., 2019, p. 20), and form 3D interconnected networks with graded molecular and cellular composition. The presence of large and small scale gradients is consistent with how tissue development is controlled (K. W. Rogers & Schier, 2011) and how epigenetics regulate cell state, but contrasts with an emphasis on enumeration of discrete cell states and mutations using single cell sequencing.

When a machine learning (kNN) model involving high-plex intensity data was trained by a pathologist to distinguish morphologies such as glandular vs. solid and high vs. low grade tumor, we found archetypal morphologies used in diagnosis were graded and intermixed to a greater or lesser degree in different specimens that did not correspond to MSI-H (hypermutant) vs. MSI-L status, suggesting that epigenetic factors rather than genetic ITH plays a dominant role. We found that differences in morphology did not map to differences in single markers, but instead to hyperdimensional features involving combinations of multiple proteins. We therefore speculate that the morphologic gradients observed in tissue specimens result from the aggregate action of several underlying molecular gradients, which may include epigenetic regulators, oncogenic signaling effectors, as well as cell-extrinsic factors such as gradients in cytokines and nutrients.

Graded changes in protein expression along tumor cell fibrils represent an interesting case in which a connection can be drawn between molecular and morphological gradients. The diagnostic criterion for a tumor bud is the presence of clusters of 1-4 cells surrounded by stroma at the tumor invasive margin (Lugli et al., 2017). Tumor buds are assumed to constitute isolated single cells or small clusters of cells with EMT-like signatures prone to infiltration and metastasis (Mani et al., 2008).

However, in agreement with an earlier H&E study (Bronsert et al., 2014), we find that buds in CRC1 are most likely to be cross-sectional views of the narrow distal tips of fibrillar structures that project from the main tumor mass. Using Delaunay triangulation to quantify these structures we find that E-cadherin and Ki-67 levels fall, with progression from the widest (proximal) to the narrowest (distal) regions of the fibrils. Delaunay triangulation identifies morphologically similar fibrils in other regions of the tumor, including as projections into the mucin network. This recurrence of morphological transitions is consistent with the idea that ITH can have a substantial non-genetic origin (Black & McGranahan, 2021; Sharma et al., 2019).

### Ensuring adequate spatial power for tissue imaging

To date, most analysis of high-plex tissue images has focused on reconstructing small neighborhoods of cells, particularly from tissue microarrays and small fields of view. However, a key technical conclusion of our paper is that even local proximity analysis is confounded by poor statistical power due to pixel-to-pixel spatial correlations that generate the structures and patterns visible in images. Whereas the number of independent samples in a set of dissociated cells (e.g., in scRNA-seq) is equal to the number of cells *(N)*, the Central Limit Theorem tells us that the effective sample size (*N*_*eff*_) for spatially correlated data in an image will always be smaller (Lavrakas, 2008). In CRCs we find that correlation length scales for biologically relevant markers can be as large ∼500 µm, making *N*_*eff*_ 100 to 1000-fold smaller than N. Thus, in many cases, TMAs and mm-scale fields of view contain only a single instance of a feature of interest, resulting in measurement error that is substantially greater than the patient-to-patient variability. This penalty is even more severe for complex properties such as neighborhood inclusion and exclusion and is sufficient to generate spurious correlations with Kaplan-Meier survival estimators.

In contrast, imaging entire slides in 2D (∼10^5^ cells) largely overcomes this problem (*N*_*eff*_ ∼ 100). It is also the standard in conventional pathology (Ghaznavi et al., 2013), and is regarded by the FDA a diagnostic necessity (Aeffner et al., 2019; Health, 2019). The argument for whole-slide imaging has not conventionally had this statistical foundation, and has instead been justified by the need to view tumor cells in context for classification using the TNM system (Amin et al., 2017), the performance of which is only rarely exceeded by the addition of molecular data. However, the two arguments are fundamentally similar. Our data also show that 3D reconstruction enables substantial additional insight into the connectivity of large-scale structures, but for relatively straightforward tasks such as cell-type enumeration, 2D whole-slide imaging appears adequate. Nonetheless, a requirement for whole-slide imaging in a research and diagnostic setting comes with substantial cost: per-patient data sets are >10^2^ larger than with TMAs, cohorts are more difficult to acquire (whole blocks must be accessed and recut), and the data analysis remains challenging since files can be as large as a terabyte per specimen.

### Immunology of the CRC invasive margin

The morphology and depth of invasion of a tumor margin has high prognostic value (Weiser, 2018) and differences between infiltrative and well-delineated pushing margins are commonly used for patient management (Koelzer & Lugli, 2014). By annotating invasive margins in our CRC cohort, we found that the immune environment can vary substantially within a single tumor and also recurrently with margin morphology across specimens. Budding regions are the most T-cell rich, but also the most immunosuppressive (with abundant T_regs_ and PDL1-expressing ells). Whereas tumor buds have few proliferating cells, tumor cells in areas of deep invasive margins are highly proliferative and have fewer immediately adjacent immune cells. Because MSI-H CRC is often treated with ICIs, the mechanism of PDL1-mediated suppression of T cells at the tumor margin is particularly relevant (André et al., 2022). We find that, in all but one of the 17 CRCs we examined, PDL1-expressing myeloid cells outnumber PDL1-expressing tumor cells 4-fold or more; high resolution imaging also demonstrates that myeloid cells form PDL1:PD1 mediated contacts with PD1^+^ T cells. These findings are consistent with recent data from mouse models of colon cancer showing that dendritic cells are a primary source of immunosuppressive PDL1 (Oh et al., 2020) and with a general role for dendritic cells in tolerization. However, we find that the relative abundance of PDL1^+^ cells most likely to correspond to macrophages and dendritic cells proximate to T cells varies substantially from tumor to tumor, suggesting that dendritic cells are not the only relevant PDL1^+^ myeloid population. Deeper molecular profiling should make it possible to determine the precise identities of PDL1^+^ myeloid cells involved in T-cell suppression in different tumors as well their prognostic significance. Although PDL1^+^ tumor cells were rare in all but one tumor (CRC17), these cells can also play a role in immunosuppression because they were often found concentrated in regions of tumor budding. An obvious follow-up question that will require analysis of cohorts of ICI-treated patients is whether the origin of PDL1 plays a role in responsiveness to ICIs and whether tumors that are exceptionally high in tumor-intrinsic PDL1 – like CRC17 – will be more or less ICI sensitive.

### Limitations of this study

The most substantial limitation in the current study is that only one CRC has been reconstructed in 3D, largely because the process remains arduous and manual. Moreover, many of the features whose architecture we examine – tumor budding fibrils, TLS networks, and invasive margins – would benefit from deeper molecular profiling to better identify cell types and states. An obvious example is to further determine the relationships between B and plasma cell maturation markers, antigen presenting cells such as dendritic cells and helper T cells, cytotoxic cells such as CD8 T cells, and functional states with regard to emergent 3D TLS morphologies. There are also many spatial relationships among the 2 × 10^8^ cells in our dataset that we are unable to explore in a single paper, particularly since several of our computational methods are quite simple (e.g., k-means clustering); other approaches from graph or percolation theory might be superior (Plotkin et al., 2002; Reynolds et al., 2009). Moreover, image segmentation and cell-type calling methods continue to improve, and all types of analysis will likely benefit in the future from reprocessing of primary images. To mitigate these and other limitations, and to enable follow-on studies, we are releasing all images and processed data in multiple formats. The results described above also suggest multiple ways in which data could be better acquired for future 3D tumor studies.

## Supporting information

Movie 2 (Budding)

Movie 3 (TLS 3D)

## DATA AVAILABILITY AND ATLAS IMAGE VIEWING (PRE-PUBLICATION)

As part of this paper all images at full resolution, all derived image data (e.g., segmentation masks) and all cell count tables have been released via the NCI-sponsored repository for Human Tumor Atlas Network (HTAN; https://htan-portal-nextjs.vercel.app/). Because the public resource is still undergoing extensive development, an additional version of the numerical data is also available at https://www.synapse.org/#!Synapse:syn18434611/wiki/597418. Several of the figure panels in this paper are available with text and audio narration for anonymous on-line browsing using MINERVA software (Rashid et al., 2022), which supports zoom, pan, and selection actions without requiring the installation of software. A Minerva story with an overview of CRC1 (sections 096 and 097) can be found at: cycif.org/crc1-intro and the 25 CRC1 Z-sections can be found at: cycif.org/crc1-3d. The third Minerva story focused on data integration for CRC1 can be found at: https://www.cycif.org/data/linwang-coy-2021/viz.html. Other resources, including images of CRC2-17, for this paper can be found at https://www.tissue-atlas.org/atlas-datasets/lin-wang-coy-2021/. We will make all of these MINERVA stories available directly via the published version of this paper; we are currently securing DOIs for these stories to provide a more uniform name space.

## ACKNOWLEDGEMENTS

We thank Alyce Chen, Raquel Arias-Camison, Zoltan Maliga, and Jeremy Muhlich for help in all stages of this project and Juliann Tefft for editing the manuscript. This work was supported by NIH grants U54-CA225088 (PKS, SS), U2C-CA233280 (PKS, SS), U2C-CA233262 (PKS, SS), U2C-CA233291 (CNH, KSL), R01-DK103831 (CNH, KSL), NIH training grant T32-GM007748 (SC), and the Ludwig Center at Harvard (PKS, SS). All HTAN consortium members are named at humantumoratlas.org. Development of computational methods was supported by the Ludwig Cancer Research, by a Team Science Grant from the Gray Foundation, and by the David Liposarcoma Research Initiative. We thank Dana-Farber/Harvard Cancer Center for the use of the Specialized Histopathology Core, which provided histopathology services supported by P30-CA06516.

## AUTHOR CONTRIBUTIONS

JRL, PKS, and SS developed the concept for the study. JRL, CY, and YC collected image data and also performed analysis in collaboration with SC and SW. CNH and KSL collected and analyzed scRNA-Seq data. JRL, SC, and MKN performed GeoMx experiments and analysis. MT developed the MINERVA stories. All authors wrote and edited the manuscript. SS, PKS, and KSL provided supervision and funding.

## DATA AND SOFTWARE AVAILABILITY

All full resolution images, derived image data (e.g., segmentation masks) and all cell count tables will be publicly released via the NCI-sponsored repository for Human Tumor Atlas Network (HTAN; https://humantumoratlas.org/) at Sage Synapse. A version of this data is available at https://www.synapse.org/#!Synapse:syn18434611/wiki/597418. Several of the figure panels in this paper are available with text and audio narration for anonymous on-line browsing using MINERVA software (Rashid et al., 2022), which supports zoom, pan and selection actions without requiring the installation of software. A Minerva story with an overview of CRC1 (sections 096 and 097) can be found at: cycif.org/crc1-intro and the 25 CRC1 Z-sections can be found at: cycif.org/crc1-3d. scRNA-seq data is available in the Gene Expression Omnibus (GEO accession: GSE166319).

All software used in this manuscript is freely available via GitHub as described in (Schapiro et al., 2022) and references therein and in https://github.com/labsyspharm/CRC_atlas_2022.

## SUPPLEMENTAL INFORMATION

Supplemental information includes six figures, eight tables, and three movies (Lin-Wang-Coy-CRC1-Movie 1-Lumen View, Lin-Wang-Coy-CRC1-Movie 2-Budding and Lin-Wang-Coy-CRC1-Movie 3-TLS). Interactive data viewing is possible via anonymous web links via https://www.cycif.org/crc1-intro, https://www.cycif.org/crc1-3d, and https://www.cycif.org/data/lin-wang-coy-2021/viz.html.

## SUPPLEMENTAL FIGURE LEGENDS

**Figure S1. Overview of dataset and connection between cell-type calling and underlying morphologies. *Related to Figure 1***.

Design of GeoMx experiment. **(A)** 32 regions were selected from one tissue section of CRC1 (31 passed quality control and were used in the analysis). Representative images of the regions are shown (right panels). Scale bars: 2 mm in left panel, 200 µm in the zoomed in views and 200 µm in the right panels showing representative morphologies. **(B)** Representative images of main antibody panel from CRC1. (Blue: DNA stain with Hoechst 33342). Scale bars, 100 µm. **(C)** Cell-types mapped across CRC1, section 097. Cell-type definitions and main classification markers are as indicated. A detailed marker/reference dictionary is presented in **Table S7. (D)** Variation in composition of each annotated ROI across all sections of CRC1 for the same three main classes of cell-types as listed in **Figure S1C** (tumor epithelium, stroma, and immune). **(E)** UMAP plot of scRNA-seq data generated from CRC1, and cell-types identified by Leiden clustering (see Methods). **(F)** Marker-guided sub-clustering was performed as described in Methods. Positive cells are highlighted in yellow.

**Figure S2. Impact of TMA sampling error. *Related to Figure 2***.

**(A)** Field of view (FOV) portraying four different correlation length scales and strengths for CK^+^, FOXP3^+^, α-SMA^+^, and CD163^+^ cells. Four circles with radii denoting the length scale parameters. **(B)** Scaling law estimates of N/N_eff_ for CK^+^, FOXP3^+^, and α-SMA^+^ based on the scaling law in Equation 1 (shaded colored boxes represent *l*_*cell*_ = 7-13 μm, *l* from fits) compared to 0.6 mm vTMA cores (colored bars). **(C)** Correlation of select cell-type pairs amongst 10 nearest neighbors. **(D)** Correlation functions between select cell-type pairs as estimated from virtual tissue microarrays (vTMAs) or random sampling, overlaid with the correlation functions from four cores (from **Figure 2I**). **(E)** Percent variance in real TMA (rTMA) estimates of cell-type abundance that can be attributed to sampling error, after removing outliers. Expected improvement from sampling four cores per tumor is shown in yellow. **(F)** FOVs of patient sections with low and high Ki-67^+^ cell abundance. Circles show the length scale of Ki-67^+^ cells. **(G, H)** Kaplan-Meier (KM) curves for progression-free survival (patients CRC2-17), calculated from TMAs (left) and whole-slide images (WSI, right). **(G)** KM curves generated from data stratified with α-SMA^+^ percentage (cutoff 40%) in each patient sample. **(H)** KM curves generated from data stratified with mean CD4 expression level (cutoff: 3,500 AFU) in each patient sample. **(I)** Variation of cell-type composition between sections of a single tumor (CRC1) and sections from different patient tumors (CRC2-17). Section sampling error is typically a minority of the variance between patient sections. **(J)** Variation of cell-type spatial correlation strengths and length scales across CRC1 Z-sections (blue) and across patients CRC2-17 (red). In most cases, variation within a patient is smaller than that between patients and shows no signs of bias.

**Figure S3. kNN-classification of epithelial histology. *Related to Figure 3***.

**(A)** Precision and recall of morphology classifiers trained on CRC1 sections. **(B)** Normalized Shannon entropy of cells in the CRC sections indicated in **Figure 3F. (C)** *(Left)* Shannon entropy of kNN-classification for cells in CRC1. Normal cells from normal colon epithelium (ROI1) have low-entropy, indicating high-confidence classification. Regions used for training were also high confidence, as expected by definition. Most tumor regions were classified as being between classes, i.e., having high entropy. *(Right)* The relative weight of each class is visualized by hue. **(D)** Dimensional reduction of subsampled single-cell expression from CK^+^ cells by t-SNE, with pathologist annotations indicated by color. Each of the four marker panels provide enough information to cluster normal epithelial cells (black) separately from tumor cells, despite limited overlap in markers between panels (indicated by Venn diagram). Different annotations roughly occupy different regions of expression space, indicating that expression and morphology are correlated, but tumor cells largely form a continuous distribution, supporting the existence of mixed morphologies.

**Figure S4. Tumor bud characterization. *Related to Figure 4***.

**(A)** Proportion of pathology annotated budding cells amongst CK^+^ cells across each of the sections. **(B)** CyCIF image with location of 3D viewpoints corresponding to **Figure 4C** and **4D**. Arrow represents approximate viewing angle in those figures. **(C)** Differential expression of markers in cells annotated as tumor buds. The relative expression of indicated markers is represented in heatmap as the log2 ratio of budding tumor cells to all tumor cells. **(D)** Log-intensity of markers and their dependency on cluster-size in CRC1 tumor cells across all 25 sections (as in **Figure 4H**). Expression of annotated buds shown in green for reference. Box plots show 1^st^-3^rd^ quartiles; points beyond are not shown. Each box represents ∼10^5^-10^6^ cells. **(E)** Log-intensity of markers and their dependency on cluster-size for tumor cells in CRC2-17 (as in **Figure 4I**).

**Figure S5. 3D TLS structures and clusters. *Related to Figure 6***.

**(A)** 2D projection of TLS networks (TLSN) across all sections of CRC1 (left panel), and the section-by-section view of TLS networks in nine selected sections (right panels). **(B)** The number of TLS per 3D TLS network (top) and the number of the total of 25 slides in which a particular TLS network was identified (bottom). **(C)** TLS domain cluster composition in each section of CRC1. **(D)** Representative H&E images of TLS domain clusters 1 and 3, the same regions as shown in **Figure 6F** (serial section shown). **(E)** 3D view of TLSN-B, colored by principal component 2 (PC2) (PC1 shown in **Figure 6H**). **(F)** t-SNE plots of all TLS domain clusters from CRC1 (25 sections) and CRC2-17 (16 sections), colored by samples (left panel) or TLS domain clusters (right panel).

**Figure S6. LDA analysis of immune composition. *Related to Figure 7***.

**(A)** 16 LDA topics from CRC1-17, immune panel (33 antibodies). Representative markers are shown in red and black text (the size of label for each marker is proportional to its probability within each of the topics). **(B)** Two-way hierarchal clustering between LDA topics and pathology annotated regions. The cell/topic counts from all pathologist-annotated ROIs as well as marker-defined ROIs (*) were clustered with full lineage and Euclidean distancing. **(C)** The fractions of topics in three selected regions of the invasive margin across all samples (a subset of the topics is shown in **Figure 7E**; budding margin, deep invasive margin, and ‘border’ margin which does not include the deepest invasive front). **(D)** Fractions of PD1^+^ cells in selected populations. The percentage of PD1^+^ cells in total/any cells or cell groups selected with indicated markers were plotted sample by sample (CRC1-17). **(E)** Fractions of PDL1^+^ cells in selected populations. The percentage of PDL1^+^ cells in total/any cells or cell groups selected with indicated markers were plotted per CRC sample. **(F)** The fractions of PDL1^+^ cells in myeloid subsets. Box plot showing the percentage of PDL1^+^ cells in CD68^+^CD11c^-^, CD68^-^CD11c^+^, and CD68^+^CD11c^+^ per sample (boxes indicate 1^st^-3^rd^ quartiles and whiskers represent 5% and 95%; red line indicates medians). **(G)** Relative frequency of PDL1+ myeloid subsets in each sample. The numbers of CD68^+^CD11c^-^, CD68^-^CD11c^+^ and CD68^+^CD11c^+^ cells in PDL1+ population were calculated, and the relative abundancy (divided by sum) of each subset is shown.

**Figure S7. Cell composition in CRC2-17. *Related to Figure 7***.

**(A)** t-SNE plots based on CyCIF data for specimens CRC1-17 (excluding data from the DNA staining). Cell types are shown at the bottom of the figure. Tumor/epithelium (T/E), stroma (S) and immune (I) populations are outlined in black. The t-SNE plot for CRC1 is reproduced from **Figure 1E** for reference. **(B)** t-SNE of CRC1-17 labelled by specimen identity with labelling by general cell type in upper right. **(C)** Cell-type composition for CRC1-17 shown as stacked bar graphs with the same color code as in panel A.

## SUPPLEMENTAL TABLE LEGENDS

**Table S1. Clinical information for colorectal cancer cohort**. Demographic and diagnostic information for all patient-derived specimens in this study. CRC1 was analyzed in 3D and CRC2-17 in whole-slide 2D and TMA 2D. Other specimens (CRC18-93) were imaged as TMA cores, as described in the text and Figure 1.

**Table S2. Sectioning plan for specimen CRC1**. Thickness and staining plan for CRC1 sections shown in Figure 1. All CyCIF sections other than 044-046 were stained using the primary CyCIF antibody panel described in Table S3. Sections 044-406 were stained as described in Table S4. Numbers refer to the HTAN universal ID scheme used to access underlying Level 2 to Level 4 data in the HTAN data portal.

**Table S3. Primary CyCIF antibody panel**. Antibodies used to stain all CRC1 sections 044-046 including CRC2-17 sections and TMAs. CST refers to Cell Signaling Technologies (Beverley MA USA); RRID refers to the Research Resource Identifier available at https://scicrunch.org/resources.

**Table S4. Supplementary CyCIF antibody panel for CRC1**. Antibodies used to stain CRC1 sections 044-046. Abbreviations as in Table S3.

**Table S5. Tumor-focused antibody panel**. Antibodies used to stain CRC2-17 for whole-slide imaging.

**Table S6. Immune-focused antibody panel**. Antibodies used to stain CRC2-17 for whole-slide imaging.

**Table S7. Cell-type dictionary**. Cell-type assignments based on marker intensities. The first tab shows the primary discriminating markers and tabs 2 and 3 show assignments based on all markers in the panel.

**Table S8. Cell-type composition for regions of interest in CRC1**. Cell-type composition for pathologist-defined regions of interest (see Figure 1B) across all sections processed for CyCIF. Cell-type definitions as in Table S7.

## STAR METHODS

### CONTACT FOR REAGENT AND RESOURCE SHARING

This manuscript contains no unique reagents or resources; all antibodies are available commercially (see **Table S3-S6** and Key Resources file) and all data can be accessed via the Human Tumor Atlas Network (HTAN) portal (https://htan-portal-nextjs.vercel.app/).

## EXPERIMENTAL MODEL AND SUBJECT DETAILS

### Human Tissue

Unfixed (fresh) tissue from a resection of a colon adenocarcinoma (CRC1) was isolated by the Cooperative Human Tissue Network (CHTN) for single cell RNA-sequencing. A portion of the sample was formalin-fixed and paraffin-embedded (FFPE) and tissue sections were generated by the CHTN as outlined in **Table S2**. Additional FFPE colon adenocarcinoma specimens were retrieved from the archives of the Department of Pathology at Brigham and Women’s Hospital (BWH) with Institutional Review Board (IRB) approval as part of a discarded/excess tissue protocol. 92 different tumor samples (CRC2-93) were used to construct a tissue microarray (HTMA 402; four 0.6 mm diameter cores were extracted from the FFPE donor blocks and assembled into a recipient TMA block). Whole-slide sections of 16 of these colon adenocarcinoma specimens (CRC2-17) were also analyzed, after the four cores were removed. Clinical metadata was abstracted from the BWH medical record and clinical and biospecimen metadata for CRC1 was provided by the CHTN. The tumor and adjacent normal tissue in CRC1 was collected from a resection of the cecum of a 69-year old male; the medical reports indicated that the tumor was a poorly differentiated stage IIIB adenocarcinoma (pT3N1bM0) (Weiser, 2018) with microsatellite instability (MSI-H) and a BRAF^V600E^ (c.1799T>A) mutation. Histopathology review showed that the tumor had a broad front invading into underlying muscle (*muscularis propria*) and connective tissue giving rise to a ‘budding margin’ (IM-A) adjacent to an area of normal colon mucosa (ROI1), a ‘mucinous margin’ in the middle of the specimen (IM-B), and a deep ‘pushing margin’ (IM-C) (these three margins are denoted “A”, “B” and “C” in **Figure 1B**).

## METHOD DETAILS

### CyCIF protocol

Tissue-based cyclic immunofluorescence (CyCIF) was performed as previously described (Lin et al., 2018). The detailed protocol is available in protocols.io (dx.doi.org/10.17504/protocols.io.bjiukkew). In brief, the BOND RX Automated IHC/ISH Stainer was used to bake FFPE slides at 60°C for 30 minutes, to dewax the sections using the Bond Dewax solution at 72°C, and for antigen retrieval using Epitope Retrieval 1 (Leica^TM^) solution at 100°C for 20 minutes. Slides underwent multiple cycles of antibody incubation, imaging, and fluorophore inactivation. All antibodies were incubated overnight at 4°C in the dark. Slides were stained with Hoechst 33342 for 10 minutes at room temperature in the dark following antibody incubation in every cycle. Coverslips were wet-mounted using 200 µL of 10% Glycerol in PBS prior to imaging. Images were acquired using a 20x objective (0.75 NA) on a CyteFinder slide scanning fluorescence microscope (RareCyte Inc. Seattle WA). Fluorophores were inactivated using a 4.5% H_2_O_2_, 24 mM NaOH/PBS solution and an LED light source for 1 hour.

### Single-cell RNA-sequencing

Samples for scRNA-seq were processed according to the HTAN publication (Chen et al., 2021). Surgical tissues were removed and placed into RPMI solution and transported directly to the processing laboratory within 10 minutes. Tissue samples were immediately minced to approximately 4 mm^2^ and washed with DPBS. The samples were then incubated in chelation buffer (4 mM EDTA, 0.5 mM DTT) at 4°C for 1□hour and 15□minutes. Then, the resulting suspensions were dissociated with cold protease and DNAse I for 25 minutes. The suspensions were triturated throughout the process, every 10 minutes, then washed three times with DPBS before encapsulation. Single cells were encapsulated and barcoded using the inDrop scRNA-seq platform as previously described (Banerjee et al., 2020), targeting about 2,500 cells. Sequencing libraries were prepared using TruDrop library structure (Southard-Smith et al., 2020). Sequencing was performed on the NovaSeq 6000 (150 bp paired end) at a depth of approximately 150 million reads per sample.

## QUANTIFICATION AND STATISTICAL ANALYSIS

### Image processing and data quantification

Image analysis was performed with the Docker-based NextFlow pipeline MCMICRO) (Schapiro et al., 2022) and with customized scripts in Python, ImageJ and MATLAB. All code is available in GitHub (https://github.com/labsyspharm/CRC_atlas_2022). Briefly, after raw images were acquired, stitching and registration of the different tiles and cycles was performed with MCMICRO using the ASHLAR module (Muhlich et al., 2022). The assembled OME.TIF files from each slide were then passed through quantification modules. For background subtraction, a rolling ball algorithm with 50-pixel radius was applied using ImageJ/Fiji. For segmentation and quantification, UNMICST2 was used (Schapiro et al., 2022; Yapp et al., 2022) supplemented by customized ImageJ scripts (Lin et al., 2018) to generate single-cell data. More details and source code can be found at www.cycif.org and as listed in the software availability section.

### Single-cell data quality control for CyCIF

Single-cell data for multiplexed images was passed through several quality control (QC) steps during generation of the cell feature table. Initial QC was done simultaneously with segmentation and quantification, so that cells lost from the specimen in the later cycles would not be included in the output. Next, single-cell data was filtered based on the mean Hoechst staining intensity across cycles; cells with coefficient of variation (CV) greater than three standard deviations from the mean were discarded as were any objects identified by segmentation as “cells” but having no DNA intensity. These steps are designed to eliminate cells in which the nuclei are not included as a result of sectioning. Highly autofluorescent (AF) cells (measured in cycle 1 or 2) were also removed from the analysis, using a customized MATLAB script that applied a Gaussian Mixture Model (GMM) to identify high-AF populations. More details and scripts are available at https://github.com/labsyspharm/CRC_atlas_2022.

### Cell-type identification using CyCIF data

Multiparameter single-cell intensity data was used for generating binary gates. For the main CyCIF panels, 16 measurements (cytokeratin, Ki-67, CD3, CD20, CD45RO, CD4 CD8a, CD68 CD163, FOXP3, PD1, PDL1, CD31, α-SMA, desmin, and CD45) were subjected to binary gating. All samples and markers were gated independently. A customized MATLAB script was used to apply 2-component Gaussian Mixture Modeling and generate the initial gate, followed by human-inspection and adjustment. Double or triple gates were also generated via Boolean operation in single-cell data. For hierarchal cell-type identification, a modified SYLARAS algorithm (Baker et al., 2020) was applied with these datasets, and a total of 21 different cell types were assigned using the 16 markers described above. Additional markers (e.g., E-cadherin) were considered to be continuous variables and used for analysis but not cell-type assignment. The completed cell dictionary for cell-type identification can be found in **Table S7**.

### Pathology annotation of histologic features

Hematoxylin and eosin (H&E) stained tissue sections from all specimens (CRC1-17) were evaluated by two board-certified pathologists (S.C., S.S.). For each case, 6 principle regions of interest (ROI) corresponding to histopathologic regions or morphologic variations defined in the pathologic evaluation of CRC were defined when present for all 22 H&E Z-levels, including: (1) normal mucosa; (2) moderately differentiated invasive adenocarcinoma (glandular, typical morphology) involving the luminal surface, (3) submucosa (corresponding to ‘pT2’ depth by TNM staging), and (4) muscularis propria (corresponding to ‘pT3’ by TNM staging); (5) poorly differentiated invasive adenocarcinoma (solid, signet ring cells, corresponding to ‘high-grade’ histology); and (6) moderately-poorly differentiated invasive adenocarcinoma with mucinous features and extracellular mucin pooling (6). Regions of ITBCC-defined tumor budding (i.e., clusters of ≤4 cells apparently detached from the main tumor mass surrounded by stroma at the tumor invasive front) were also annotated in CRC2-17 and on all 22 H&E Z-levels of CRC1. For CRC2-17, additional histologic features that were not present in CRC1 were also annotated when present, including: adenoma (tubular), tumor necrosis, comedo necrosis, squamoid, pleomorphic, and extensive signet ring cell tumor morphology, and perineural or lymphovascular invasion by tumor. In cases with clear anatomic orientation, the deep invasive tumor front was initially delineated as a band with an approximate width of 5-10 cell diameters (50-100 μm) at the deep edge of the tumor. In cases with multiple histologic subtypes present at the invasion margin, each type was annotated separately; in CRC1, this included IM-A (budding/infiltrative), IM-B (mucinous), and IM-C (pushing) margins, with similar notation used in other cases. Tertiary lymphoid structures were defined in each case by identifying aggregates of lymphoid cells on H&E and correlating with CD20, CD4, and CD8 immunofluorescence (CyCIF) to identify discrete aggregates of B cells with adjacent or intermixed T-cell populations, including both immature/early TLS without histologic evidence of well-formed germinal centers, and more mature TLS with germinal center formation (Fridman et al., 2022).

### Pathologist-annotated budding cells and Delaunay cluster-sizes of cytokeratin^+^ cells

Using ITBCC criteria, a trained pathologist annotated budding regions in CRC1 (n = 25) and CRC2-17 (n = 16) from both CyCIF and H&E images. These selected ROIs were used in the data analysis, and CK^+^ cells in these areas were labelled as “budding tumor cells.” In cluster size analyses, a neighborhood graph was constructed for all segmented cell centroids using Delaunay triangulation, removing edges whose lengths were greater than 20 μm. Then, the CK^+^ neighborhood graph was defined as the subgraph restricted to the CK^+^ cells (i.e., removing all nodes and edges connected to CK^-^ cells). The cluster size of each CK^+^ cell was defined as the number of nodes in its connected component of the subgraph. For quantification of marker expression dependence on cluster-size, cells annotated as normal colon mucosa (ROI1) were removed from the CK^+^ subgraph. In the 25 CRC1 Z-sections, cells in the upper-left corner of the image (1 cm x 1 cm) were also removed; this region contained CK^+^ cells of reactive, benign, and mesothelial origin, as opposed to tumor cells of interest.

### Biased downsampling based on cluster-size for t-SNE visualization

By definition, most tumor cells have a large cluster-size. Therefore, to visualize the cluster-size dependence of marker expression with t-SNE, we downsampled cells by stochastically rejecting cells at frequency 1 − (1/*n*_*c*_)^4^, for cluster-size *n*_*c*_. The power of 4 was chosen empirically to balance the representation of various cluster sizes. Final t-SNE plots were made by further subsampling 1,000 cells from each section uniformly. The t-SNE plots in **Figure 4G** were computed using the following markers: Na-K ATPase, Ki-67, cytokeratin, PDL1, E-cadherin, vimentin, CDX2, lamin ABC, desmin, and PCNA.

### kNN-classification of epithelial cell morphologies trained on pathologist annotations

To develop a kNN classifier for pathologist-annotated regions of interest (ROIs), epithelial cells were defined by gating using a univariate, 2-component Gaussian Mixture Model on the relevant marker (cytokeratin, cytokeratin 19, cytokeratin 18, or E-cadherin) in each section. A kNN-classifier was trained on the annotated, epithelial cells using CyCIF marker expression as predictors, and annotated ROI labels as responses. Markers that exhibited unexpected optical artefacts or significant tissue loss were not used (see below for specific markers that were excluded). Learning and prediction were performed using MATLAB’s *fitcknn*() and *predict*() functions, with *k* = 40 neighbors. The prior probability of each label was set as uniform. In each section, there were at least 2,000 annotated cells for each label. Annotated cells were split 50/50 into training and validation sets. Posterior probability colors in **Figure S3C** (panels in right column) were visualized based on its vector of classification posterior Probabilities (*p*_1_, *p*_2_, *p*_3_, *p*_4_,), for 1: normal, 2: glandular classes, 3: solid, and 4: mucinous. The RGB-values of each cell were then defined as:

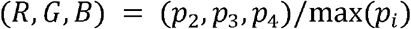

to capture the relative weight of each class.

For the sections in the primary CRC1 dataset (e.g., section 044), the following markers were used as predictors: Na-K ATPase, Ki-67, keratin, PDL1, E-cadherin, vimentin, CDX2, lamin, desmin, PCNA, autofluorescence; see paragraph below for further details on included and excluded markers. For CRC1 section 046, which was stained with an extended antibody panel, the following markers were used as predictors: cyclin B1, cytokeratin 20, cytokeratin 18, NUP98, cytokeratin 8, PDL1, acetyl-tubulin, p62, pan-cytokeratin, lamin A/C, tubulin. For sections CRC1 sections 045 and 047, which were also stained with different extended antibody panels, we used all artefact-free markers (totaling 29 and 36 respectively). For CRC2-17, the entire antibody panel was used.

In the primary dataset, for kNN classification we excluded Hoechst, CD3, CD4, CD20, CD163, CD45, CD68, FOXP3, CD45RO, α-SMA, PD1, CD8a, CD31, collagen, and autofluorescence as being irrelevant to tumor-intrinsic feature expression. The Ki-67 (D3B5) Rabbit mAb was included because it showed superior staining to another Ki67 antibody (Ki67_570) which was excluded. For CRC1 section 045, we excluded Hoechst and autofluorescence. CK17 was excluded due to staining artefacts. CK14, alternate pERK, Cyclin B1, Perforin, MAP2, GFAP, Cyclin A2, p-mTOR, Cyclin E were excluded due to tissue loss in the final cycles. For CRC1 section 046, we excluded Hoechst, autofluorescence, CD3, CD4, CD57, CD163, IBA1, CD16, CD11c, CD45, CD68, CD11b, CD11a, CD1a, Granzyme B, CD14, PD-1, HLA-A, CD8a, and CD31 as irrelevant to tumor extrinsic programs. PAX5, POLR2A, NFATc1, PAX8, and phospho-BTK were excluded due to tissue loss in late cycles. VEGFR2 was excluded due to the presence of staining artefacts. For CRC1 section 047, we excluded Hoechst, autofluorescence, and CD20 as irrelevant to tumor expression. EZH2, phospho-CDK, E2F1, FOXA2 were excluded due to staining artefacts.

### Contour plots of epithelial cell marker expression gradients

Contours represent level sets for the average marker expression of the 400 nearest tumor cells, and were computed using the MATLAB *contour()* function.

### 3D registration of CRC1 serial sections

All CyCIF sections were registered using a custom script written in MATLAB 2018 (MathWorks). Briefly, each section was first registered using a rigid transformation followed by elastic deformations starting at section 012 and cascading towards the top and bottom sections. For the rigid transformation, an early cycle Hoechst signal with minimal artefacts from each section was selected. All channels were padded by an equivalent of 1,600 pixels along all borders when registering at full resolution. Rigid transformation required consistent landmarks across all sections. Therefore, we identified two such features: the edge of the mucosa section and a point where it transitions into the stromal region. This region was annotated on several downsampled sections, providing training data for a UNet model to estimate fuzzy locations of the transition point and the mucosal edge. Starting from section 012 and taking the centroid of each fuzzy estimate as that section’s transition point, all 25 sections were aligned by translation. Each section was then rotated around the transition point until the fuzzy estimates for the edge of the mucosa region overlapped maximally between sections. For subsequent elastic deformation, we manually selected between 25-35 control points across each section. Most control points were located near the site of budding cells. Then, using local weighted means with these control points via the *fitgeotrans*() MATLAB function, we applied a deformation starting from section 012 towards section 001 and 025. Finally, we applied Demon’s algorithm to refine registration further. Images were downsampled by a factor of 0.25 and histogram matched, before applying the *imregdemons*() MATLAB function with an accumulated field smoothing of 1.5 and downsampling with 7 pyramid levels. Demon’s algorithm was applied starting from section 12.

### 3D visualization of registered CRC1 serial sections

Using Imaris, images were Gaussian-blurred, and an intensity threshold was applied to define regions (e.g., CK^+^). Connectivity of buds or mucin pools were defined on blurred, thresholded voxels.

### Virtual TMA cores and fold-change in effective sample size N/N_eff_

Virtual TMAs (vTMA) were constructed from whole-slide sections by randomly selecting a central cell and including all cells within 500 μm of the central cell’s centroid as one core. For each vTMA core, a matching, uniform random sample was generated from the whole-slide section with an equal number of cells. The standard-errors of the mean from vTMA (i.e., regional) sampling (*σ*_*TMA*_) or random sampling (*σ*_*random*_) were estimated from the means of 1,000 cores and their matched, random samples. The effective sample size N/N_eff_ was defined as the square of the standard-errors’ ratios:

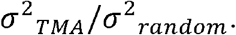

### Spatial correlation functions and predicting standard-error of regional sampling

For each sample (whole-slide, virtual TMA core, or real TMA core), spatial correlation functions *C*_*AB*_(*r*) were calculated for a pair of variables *A, B* and a nearest-neighbor index *r*. Specifically, *C*_*AB*_(*r*) was given by the Pearson correlation between cells’ *A*-values and their *r*^th^ - nearest neighbors’ *B*-values. Each *r* index was associated to the average, inter-cell-centroid distance *d*(*r*) of all *r*^th^ - nearest neighbors in a sample. Correlations were computed up to *r* =200.. Each *C*_*AB*_((d*r*)) was fit to an exponential *C*_*0*_exp (*c*_1_*d*) for parameters *C*_0_, *C*_1_, over the range of 5 < *r* < 200 to avoid spurious correlations between adjacent cells that may arise from image segmentation errors. Correlation strength was defined as *C*_0_, and length scale *l* = − 1/ *C*_1_. Fits were performed with the fit() MATLAB function with default options. We subsequently estimated the standard-error of the mean of a variable *A* for a regional sample of *N* correlated cells as follows. First, we computed the *N × N* matrix of inter-cellular distances *d*_*ij*_, and then computed the *N × N* correlation matrix *Σ*_*N*_ between cells using the fit of the spatial correlation function *C*_*AA*_(*d*). By the Central Limit Theorem for weakly-dependent variables (Ibragimov, 1962), we expect the standard-error of the mean for N samples to be 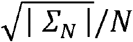,for | *Σ*_*N*_ | the sum of all entries in *Σ*_*N*_.

### Scaling analysis of fold-change in effective sample size N/N_eff_

For a variable *A* with variance *σ*^2^ = 1, the fold-change N/N_eff_ is defined as:

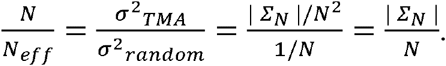

The final term can be interpreted as the sum of correlations between an average cell and all other cells in the sample region *R*. Choosing a coordinate system with an average cell at the origin, we approximate the sum as an integral:

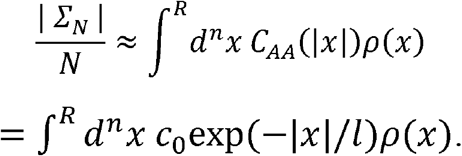

Where *ρ*(*x*) is the density of cells, and is the spatial dimension of the regional sample. If we assume a uniform density *ρ*(*x*) ∼ 1/*l*^*n*^_*cell*_ for a cell length scale *l*_*cell*_, and change variables in the integral to eliminate the length scale *l*, we have:

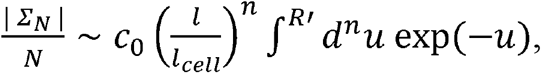

which gives us a scaling relation with which we can roughly estimate N/N_eff_ from parameters:

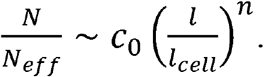

### Variance between patient TMAs due to sampling error and an optimal score

For any given cell-type’s %-composition, we computed the variance of estimates from the whole-slide tumor regions of each patient, *σ*^2^_*patient*_, and the variance of estimates from TMA cores, *σ*^2^_*TMA*_. We considered *σ*^2^_*patient*_ to be the biological variance of *σ*^2^_*TMA*_, and remaining variance to be residual error from sampling, *σ*^2^_*sampling*_. Percent of variance explained by sampling was given by *σ*^2^_*sampling*_/*σ*^2^_*TMA*_. For the hypothetical scenario of averaging 4 cores, *σ*^2^_*sampling*_ would be 4-fold lower, and percent variance explained was given by (*σ*^2^_*sampling*_/4)/ (*σ*^2^_*sampling*_/4 + (*σ*^2^_*sampling*_). Outliers in each distribution, as indicated in each boxplot, were excluded from the variance calculations.

### Immune profiling, LDA analysis, and PDL1:PD1 interaction

For CRC1-17 whole-slide sections stained with the immune panel, multiparameter single-cell intensity data was used to generate binary gates (for 30 of 33 markers). LDA analysis for spatial topic analysis was performed using MATLAB “fitlda” function. In brief, the single-cell data of each sample was split into 200 microns x 200 microns grids, and the positive frequency for each marker was calculated for each grid. The pooled frequencies of all samples were used to train the final LDA model, and 16 topics were isolated. To determine PDL1:PD1 interactions in single-cell data, the cell neighbors within 20 microns were identified with a k-nearest searching algorithm. The PDL1^+^ cells with PD1^+^ cells in proximity were labeled as “PD1^+^ interactors.” The marker expression of PD1^+^ interactors and other PDL1^+^ cells were compared as described. In **Figure 7F** (top panel), number PDL1^+^ cells with indicated subsets (any, CK^+^, CD68^+^, and CD11c^+^) were divided by the total cell number in the given subset. In **Figures 7I** and **7J**, the positive ratios were calculated by the positive cell number of indicated markers (CK^+^, CD45^+^, HLA-A^+^, and CD44^+^) normalized with the PDL1^+^ cells in either interacting or non-interacting groups.

### scRNA-seq data analysis

Following sample demultiplexing from the sequencer, reads were filtered, sorted by their barcode of origin, and aligned to the reference transcriptome to generate a counts matrix using the DropEst pipeline (Petukhov et al., 2018). Barcodes containing cells were identified using dropkick (Heiser et al., 2020). Batches were combined and consensus non-negative matrix factorization (cNMF; (Kotliar et al., 2019)) was performed to identify metagenes in the resulting cell matrix, assigning “usage” scores for each factor to all cells. The factors or metagenes contain gene loadings that rank detected genes by their contribution to each factor, which are shown on UMAP embeddings in descending order. CytoTRACE (Gulati et al., 2020) was also run using the web portal at https://cytotrace.stanford.edu/ to calculate “stemness” or cellular plasticity scores based on genetic diversity. Leiden clustering (Traag et al., 2019) and PAGA (Wolf et al., 2019) graph construction was performed on principal component analysis of the normalized and arcsinh-transformed raw counts matrix. A two-dimensional UMAP (McInnes et al., 2020) embedding was then generated using SCANPY (Wolf et al., 2018) based on principal component analysis and initial cluster positions determined by PAGA.

### GeoMx RNA spatial transcriptomics

We used the GeoMx® Cancer Transcriptome Atlas (CTA) to profile RNA expression levels of ∼1,800 genes from 32 selected regions (**Figure S1A**) from an FFPE tissue section of CRC1 using methods described by the manufacturer (NanoString Technologies, Seattle, WA). Probes were collected separately from CK^+^ and CK^-^ cells and processed using cDNA library preparation methods. The library was then sent for sequencing with Illumina NovaSeq 6000. QC was performed using vendor-provided software. 31 of the 32 samples passed QC, and these datasets were used for downstream analysis. Probe counts were normalized with the total counts in each condition and used for principal component analysis and hierarchical clustering.

