## Supplementary figures and images for "Multiplexed 3D atlas of state transitions and immune interactions in colorectal cancer"

### Supplementary Figures

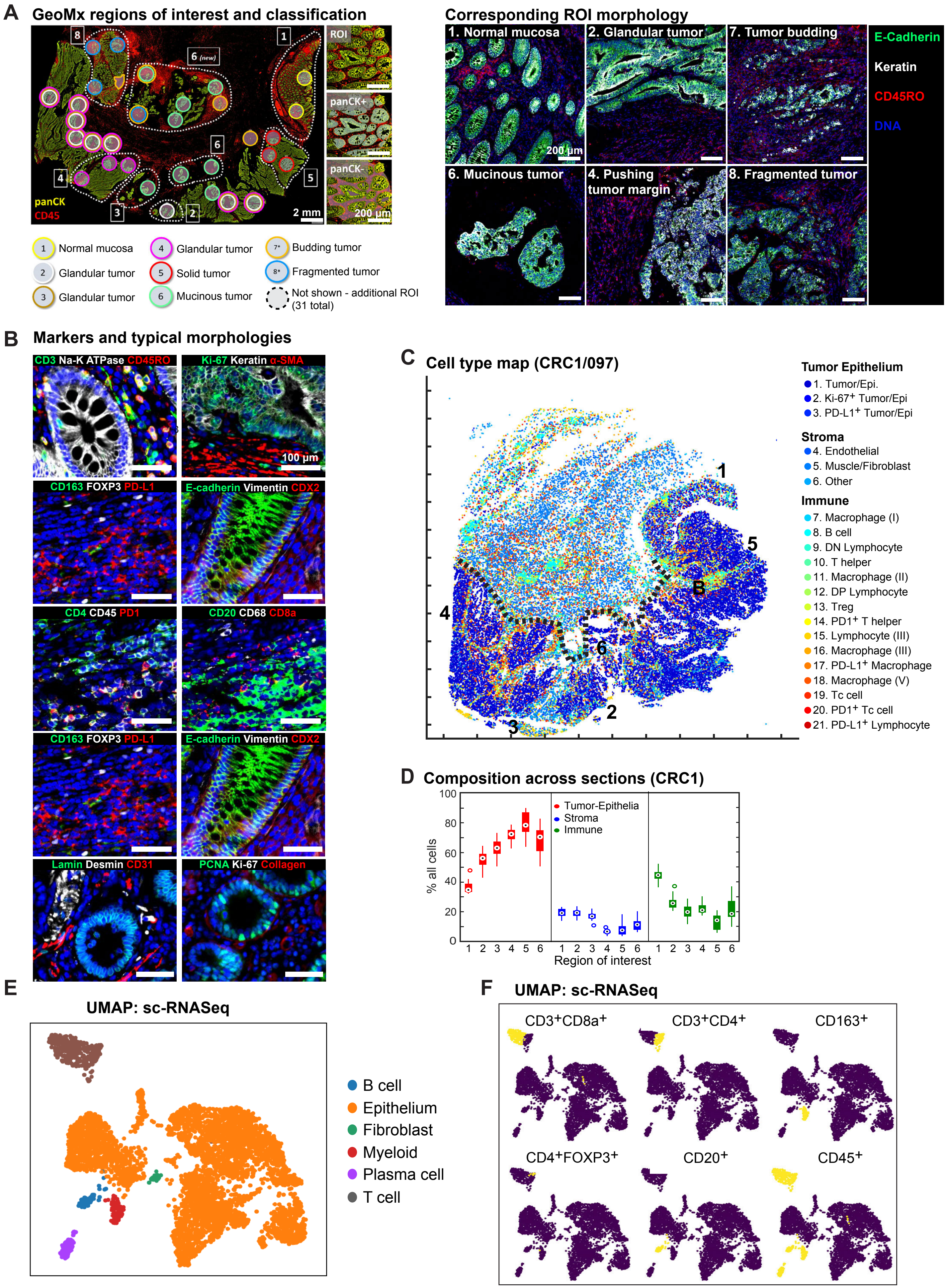

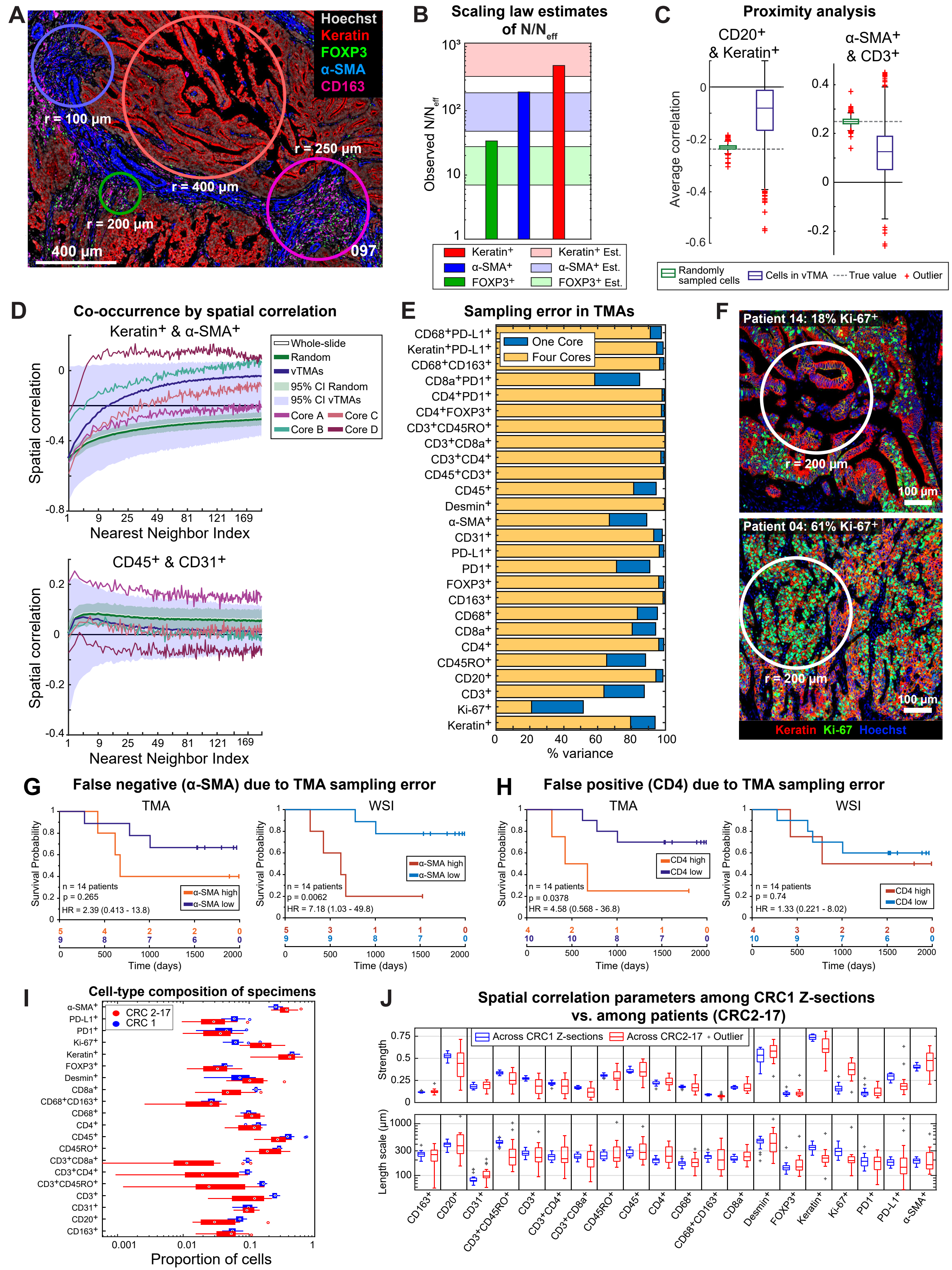

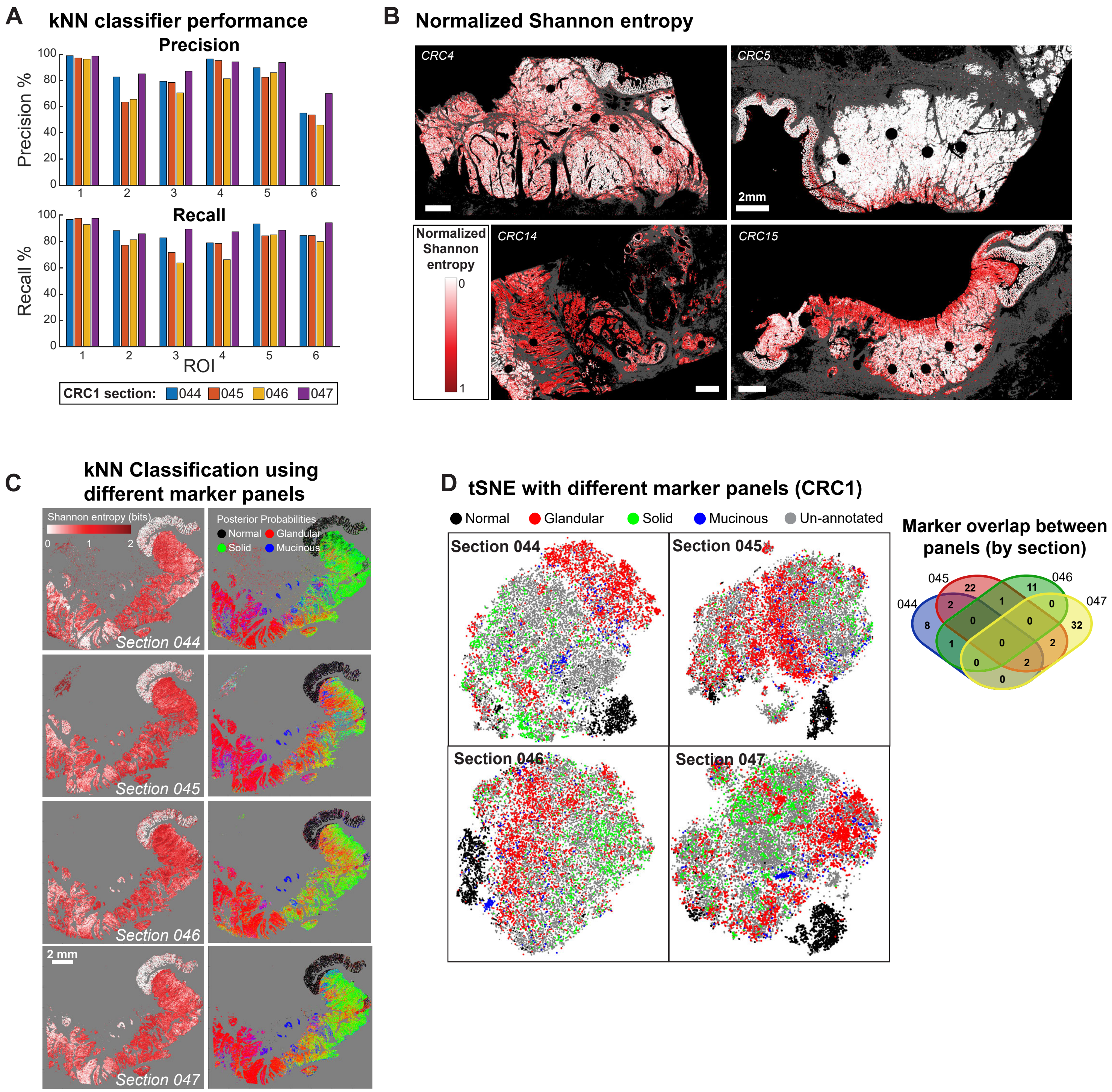

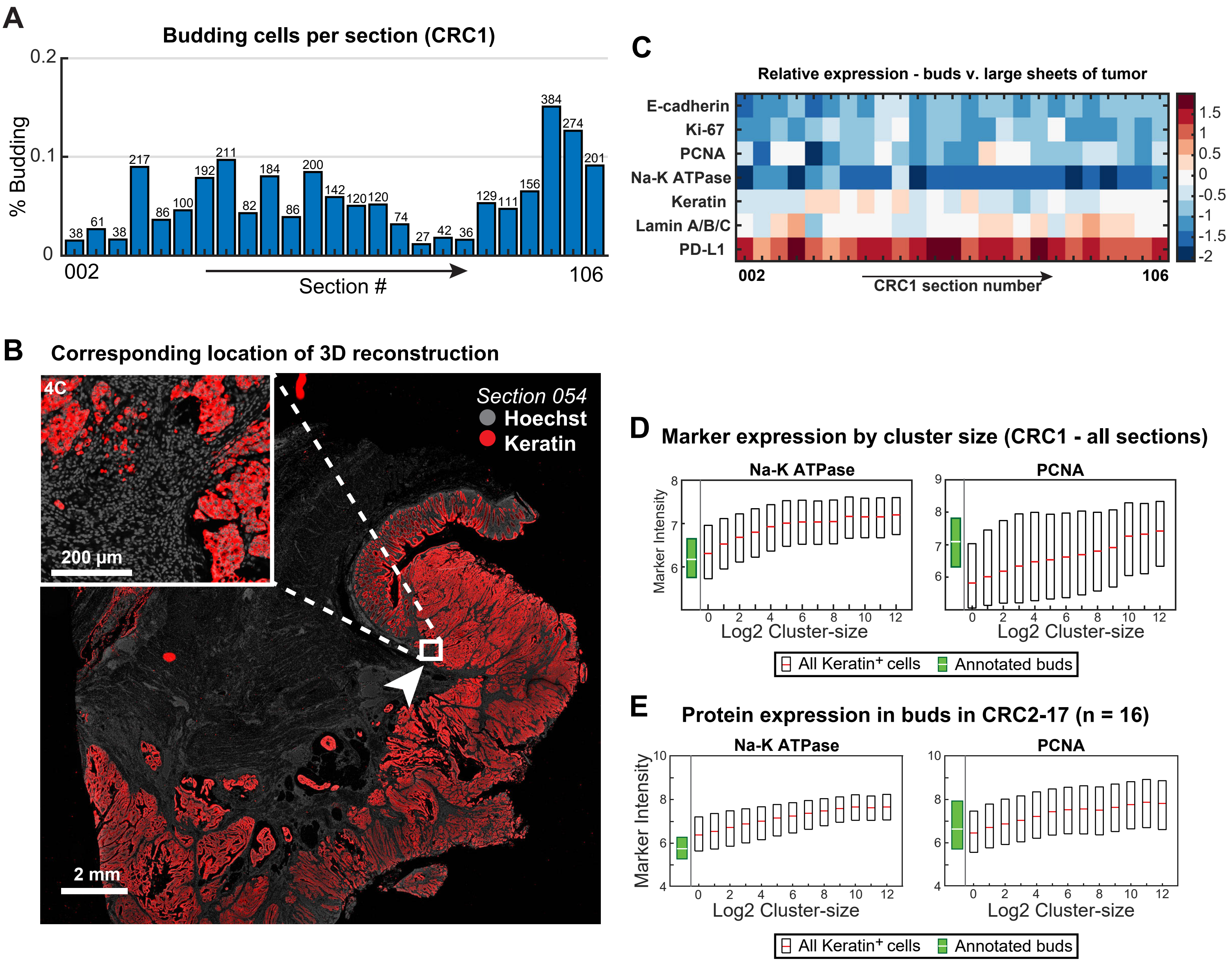

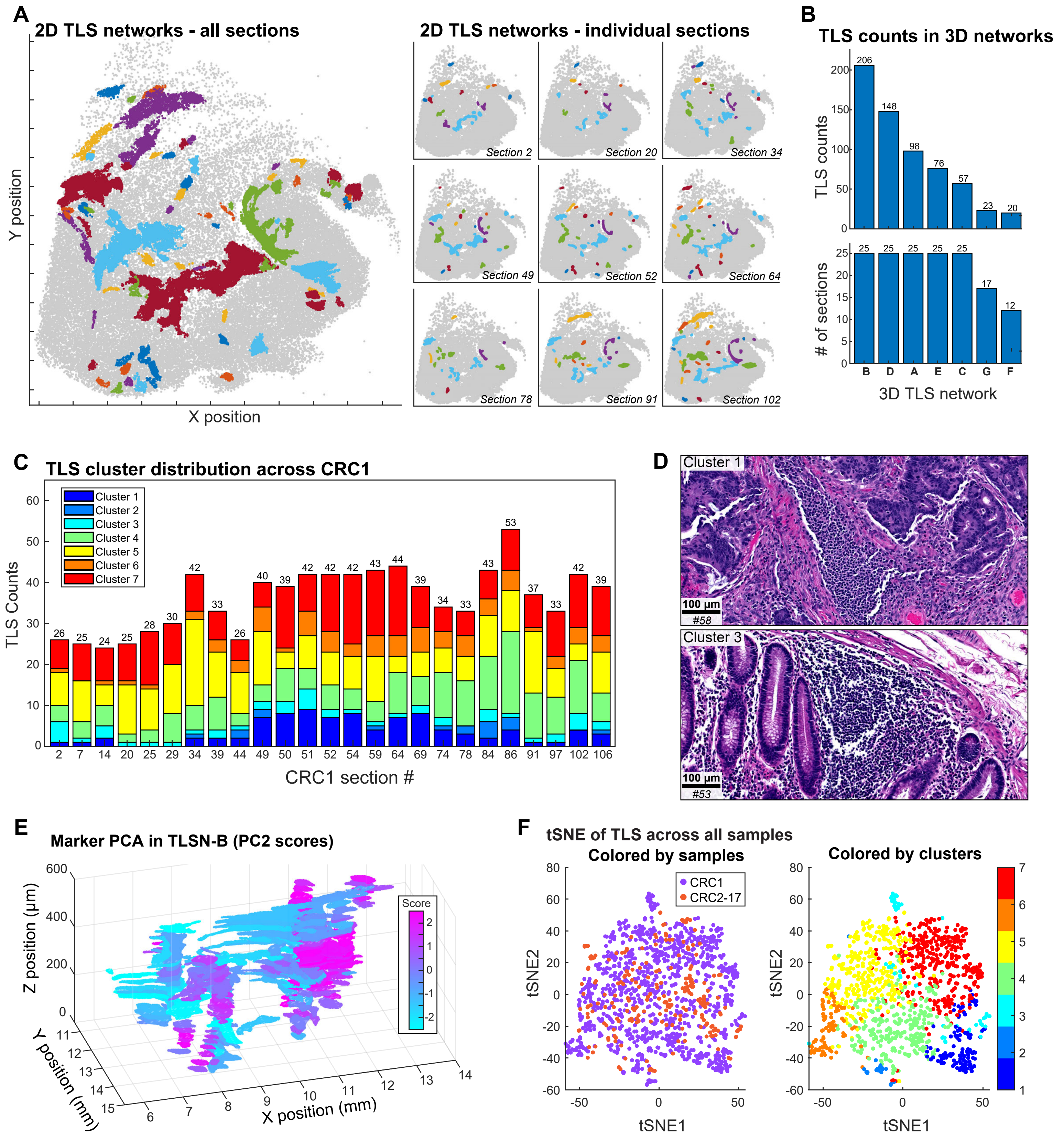

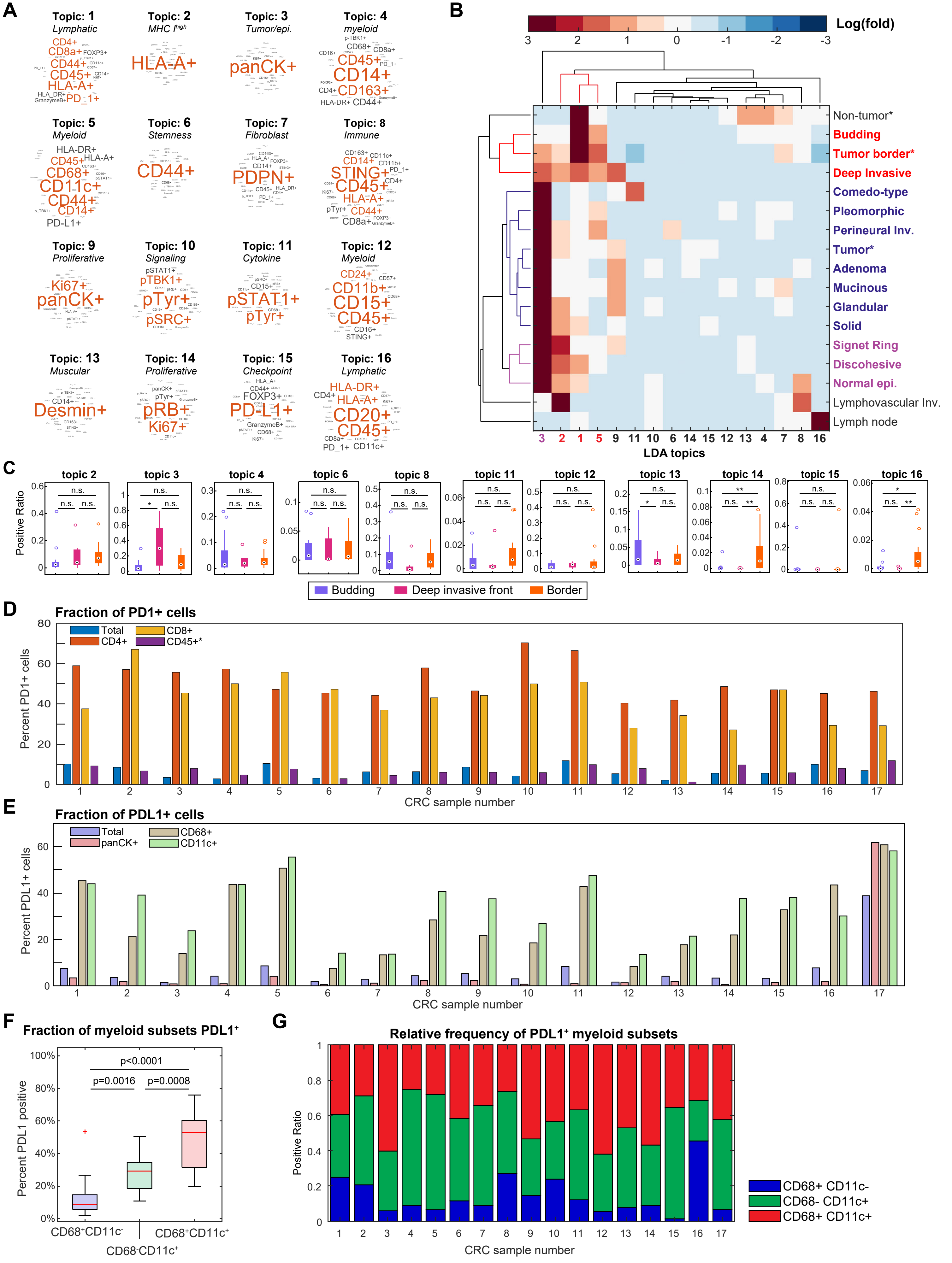

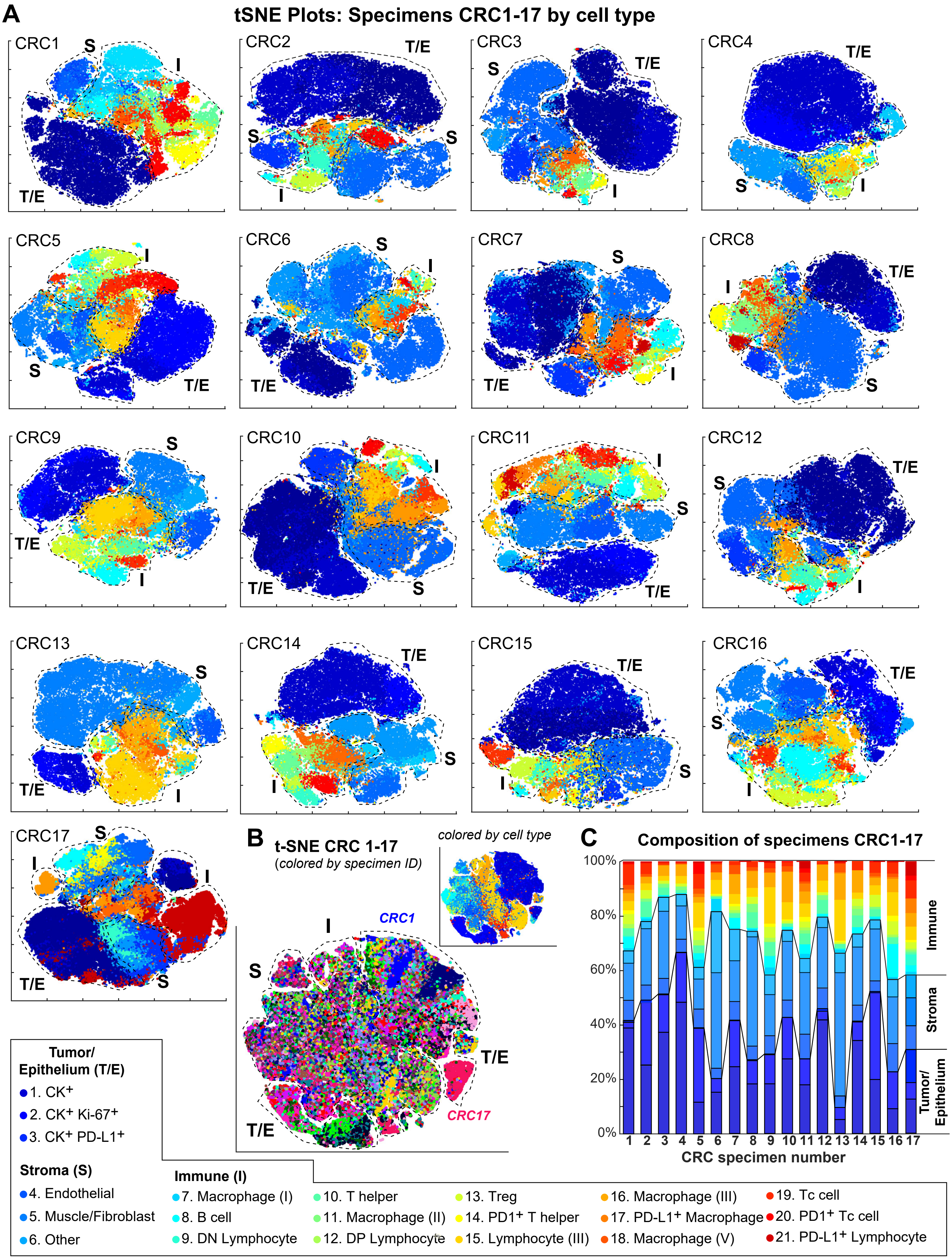
